# An Automated Scientist to Design and Optimize Microbial Strains for the Industrial Production of Small Molecules

**DOI:** 10.1101/2023.01.03.521657

**Authors:** Amoolya H. Singh, Benjamin B. Kaufmann-Malaga, Joshua A. Lerman, Daniel P. Dougherty, Yang Zhang, Alexander L. Kilbo, Erin H. Wilson, Chiam Yu Ng, Onur Erbilgin, Kate A. Curran, Christopher D. Reeves, John E. Hung, Simone Mantovani, Zachary A. King, Marites J. Ayson, Judith R. Denery, Chia-Wei Lu, Phillip Norton, Carol Tran, Darren M. Platt, Joel R. Cherry, Sunil S. Chandran, Adam L. Meadows

## Abstract

Engineering microbes to synthesize molecules of societal value has historically been a time consuming and artisanal process, with the synthesis of each new non-native molecule typically warranting its own separate publication. Because most microbial strain engineering efforts leverage a finite number of common metabolic engineering design tactics, we reasoned that automating these design steps would help create a pipeline that can quickly, cheaply, and reliably generate so-called microbial factories. In this work we describe the design and implementation of a computational system, an Automated Scientist we call Lila, which handles all metabolic engineering design and optimization through the design-build-test-learn (DBTL) paradigm. Lila generates metabolic routes, identifies relevant genetic elements for perturbation, and specifies the design and re-design of microbial strains in a matter of seconds to minutes. Strains specified by Lila are then built and subsequently phenotyped as part of a largely automated in-house pipeline. Humans remain in-the-loop to curate choices made by the system, helping for example to refine the metabolic model or suggest custom protein modifications. Lila attempted to build strains that could produce 454 biochemically diverse molecules with precursors located broadly throughout the metabolism of two microbial hosts, *Saccharomyces cerevisiae* and *Escherichia coli*. Notably, we observed the highest published titers for the molecule naringenin, the metabolic precursor to flavonoids. In total we created hundreds of thousands of microbial strains capable of overproducing 242 molecules, of which 180 are not native to *S. cerevisiae* or *E. coli*.

## Introduction

Many public and private organizations commonly use industrial fermentation of engineered microbes to make commercially viable or societally valuable products. These products, either a pure molecule or a mixture of molecules, include pharmaceutical ingredients^1^, vitamins^2–5^, flavors & fragrances^4,6^, pesticide alternatives^7^, plastics^8,9^ and polymers^10^, lubricants^11^, and dozens of others that would otherwise come from petrochemical processing, endangered plant species, or an expensive or volatile supply chain. Although industrial fermentation of engineered microbes has proven impactful in that it can deliver high purity biomolecules in sustainable fashion at stable cost, the R&D process of bringing a single biomolecule from idea to small scale production (milligrams) to applications testing (kilograms) to market (tons) is lengthy (years to a decade or more) and expensive ($100-$200M), a barrier that prevents many new biomolecules from going to market^12^.

This R&D process involves several stages of differing costs and durations. The first step involves identifying a molecule of interest. This may last from weeks to months, with a focus on market research, estimated production and purification cost, and technical feasibility. Once a molecule has been identified, the next step is to delineate the biosynthetic pathway. This is typically done manually by a trained metabolic engineer, who must either identify candidate pathways from literature or derive *de novo* pathways using expert knowledge of what biochemical reactions have precedent. Once the biosynthetic pathway has been delineated, any required enzymes that are not present in the host species must be identified, codon-optimized for expression in the host, ordered for gene synthesis, and prepared for insertion into the host. This step typically takes an additional 1-2 weeks of literature search, phylogenetic analysis, modification (for example, to remove unwanted intracellular targeting sequences), and submission of a gene synthesis order. Once the synthesized genes have been delivered, the DNA constructs must be assembled into a pathway *in vivo*, its activity confirmed, and the target molecule detected. Depending on the number of iterations required to synthesize the product, these steps take anywhere from a few months to a year. Once a proof-of-concept strain has been created that can produce detectable amounts of the molecule in the milligram range, various facets of cellular metabolism and physiology must be optimized until the strain is capable of producing the molecule at titers in the 0.5 - 5 g/L scale. This phase may include balancing enzyme co-factors, improving product export, overcoming enzymatic bottlenecks, and improving cell health and genetic stability. This optimization step tends to be the most time-consuming phase and can vary from years to a decade or more, depending on the product^12^. The next phase is to continue the optimization while starting pilot scale fermentations to produce and purify kilogram-levels of the molecule that will allow its testing for various applications. While this testing is ongoing, strain optimization continues until a cost target is finally reached that enables the scale up of production, typically another few months to a year.

Strain optimization tends to be the rate-limiting step in the R&D process both in time and cost because a vast biological search space must be explored^12^ by either rational design of a desired genotype & phenotype, or random mutagenesis followed by screening for a desired phenotype. For instance, in the rational approach to strain engineering^13^, there may be four potential biosynthetic routes to the target molecule starting from the carbon source (in our case sugar). Each of those routes are frequently composed of twenty or more biochemical reactions, some of which may be native to the organism and require tuning or changes to regulation, whereas others may be non-native and require the expression of heterologous genes. Each reaction may be catalyzed by hundreds of different possible enzymes. Certain enzymes may have dozens of subunit choices, with half a dozen choices for where to truncate an enzyme for optimal expression, and dozens of potential codon optimizations for optimal translation. Other enzymes, such as P450s, may require helper proteins to become active. Each enzyme may have hundreds of loci and promoter-terminator permutations in which to insert into the host genome for optimal pathway balancing. Thus, the total design space accessible to a typical strain engineer is the combinatorial space of these factors, which must be combined carefully, as factors can be fully crossed (any engineered protein could be expressed with any validated promoter), partially crossed (any protein could be integrated at any genomic locus unless it would create a loop out), or nested (each P450 may have only a few cognate reductase candidates). Together the total design space of potential strains exceeds ∼10^24^. The classical approach to strain improvement^14^, which entails genome-wide or site-directed mutagenesis followed by screening for a desirable trait, is invaluable when the targets for optimization are not known *a priori*. In this case enumerating the number of non-silent base pair changes in a 12MB genome results in ∼10^30^ possible mutants. With such a vast search space, it can take hundreds of iterations of DBTL cycles to optimize the production of a molecule from milligrams to kilograms. Assuming that each DBTL cycle takes weeks to months, scaling molecule production from milligrams to kilograms can take years.

One way to reduce the risk of the R&D process is to work on multiple similar molecules at once, expecting that a few will successfully scale. To do this cost-effectively, however, requires some level of automation so that each new molecule added to the R&D portfolio does not linearly add cost in the form of time or headcount. This cost reduction can be achieved if the multiple molecules are deliberately chosen to be biosynthetically and chemically similar^15^. A second way to reduce the risk of the R&D process is to diversify the kinds of molecules in the pipeline, expecting that a few will successfully commercialize. This is more challenging in practice, as the automation and analytical chemistry may not easily generalize to dissimilar molecules (e.g. flavonoids vs. terpenes), and the strain designs cannot be shared across molecules without additional cost in the form of time or headcount.

In this paper, we describe the design and implementation of an Automated Scientist software platform we named Lila that mitigates these two risks: increase the number of molecules under development, and increase the diversity of molecules under development, without significant manual intervention. Lila combines mathematical modeling and design principles generally adopted by scientists trained in the field to match a genotypic hypothesis or experimental design to the resulting phenotypic data.

The notion of automating various aspects of microbial synthesis, or even laboratory protocols, is not new. Various groups have successfully demonstrated the computational representation of biochemical reactions^16,17,18,19,20^, automation of biosynthetic route finding^21,22,23^, and laboratory protocols^24,25,26,27,28,29,30^. In the area of computational chemistry, the idea of automated synthesis is also gaining traction^31,32,33,34,35^.

As impressive as these previous efforts have been, several factors set our work apart from previous efforts. Most important is scale. We have attempted more molecules in parallel than has previously been reported by at least an order of magnitude. We also are the first to report full industrialization, in terms of the development of high-throughput pipelines to undertake design, build, test, and learn phases with near-touchless handoffs. and rate of molecule optimization (in terms of the time elapsed between first detection of a molecule and its scale-up to kilogram amounts). With the Lila platform we have shortened the timeline for molecule proof-of-concept production (i.e., zero to detectable) from months to weeks, decreased the cost of molecule development to <$10M per molecule (10-fold reduction), and simultaneously handled several hundred molecules in a multiplexed pipeline. Over the course of 105 partially overlapping DBTL cycles, Lila made >100,000 *in silico* designs targeting the production of 454 small molecules (Figure 1), ordered 1,850 genes for synthesis selected entirely by algorithm, created 32,000 distinct microbial strains (involving laboratory operations that assembled 280 million bps of DNA and transformed ∼180 million bps of DNA), and analyzed more than 10,000,000 data points. Taken together, we achieved a hit design in half of the molecules targeted and established a reproducible and largely automated process for biosynthesizing almost any small organic molecule in a microbial host.

**Figure 1.**
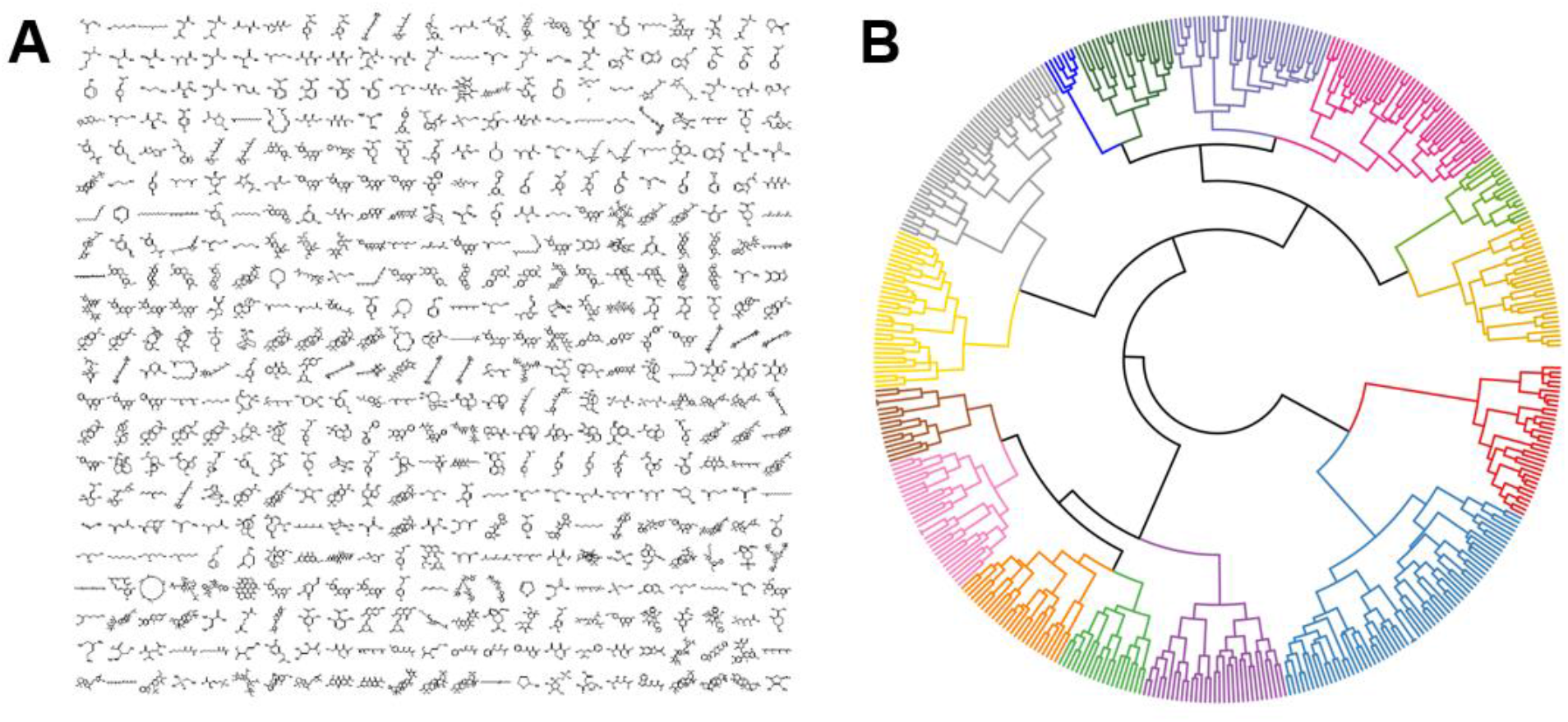
(A) Structures of molecules targeted by the Automated Scientist. (B) Chemical dendrogram of 454 molecules whose biosynthesis was attempted by the Automated Scientist, illustrating the structural diversity of molecules into 15 major classes. To generate this clustering, we retrieved Structure Data Files from PubChem for each molecule and used atom pair similarity (Tanimoto distance) to calculate a distance metric between molecules. We then hierarchically clustered the distance matrix of Tanimoto coefficients with Ward’s linkage.

## Results

### An Automated Scientist has modules akin to the cognitive functions of a human scientist

The Lila Automated Scientist software is broken into subtasks or cognitive functions that a human scientist would employ to approach a complex problem. Just as a human scientist perceives new information, learns, remembers, thinks, and executes, so too there are Lila software modules corresponding to memory, perception, reasoning, and execution. Lila was initialized with rules known to experts in the field in the Route Finding and Strain Designer modules, which interact with the Knowledge Store database that stores genotype and phenotype data. These data are used by a Strain Designer module analogous to the cognitive function of a human scientist that creates abstract genetic design rules, and decides among which of several strain improvement strategies (enzyme improvement, mutagenesis, rational engineering, or deconvolution of mutations) to pursue next. The abstract design rules are compiled into a detailed DNA sequence implementation by the Genotype Generator module, analogous to the execution function of a human scientist. Finally, the Phenotype Analyzer and Process Analyzer modules, which implement the perception and learning functions of a human scientist, analyze multivariate data and reason about uncertainty. All these modules work together with the rest of the Amyris pipeline to select molecules and analyze strains. These modules are related as shown in Figure 2.

**Figure 2.**
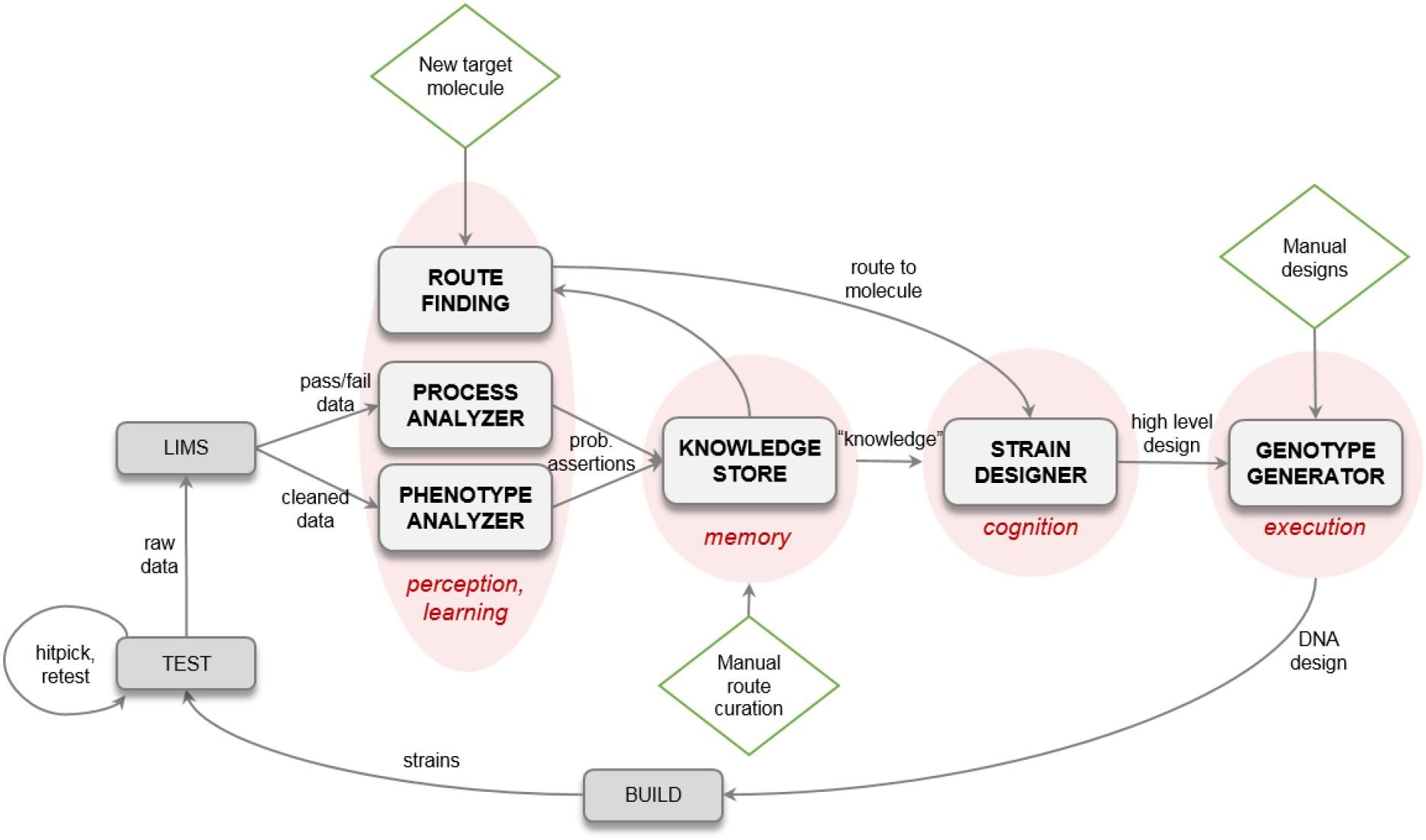
Parts of the Lila platform are analogous to a human scientist’s workflow. Green diamonds represent input points in which human scientists interact with Lila; “prob. assertions” are probabilistic assertions, i.e. scientific hypotheses with associated probabilities

More concretely, to go from target molecule to physical strain design the Lila platform makes decisions at several levels of abstraction (Figure 3). The lowest level of abstraction (called the Molecules layer) is a set of decisions around what molecules to include within the model, and how they are represented *in silico*. For example, intracellular metabolites that are transient as a reaction proceeds may be excluded to simplify a model. Next the feasible biochemical reactions within the chosen host organism (Reactions layer) are encoded and associated with physical properties such as their Gibbs free energy. These reactions are then aggregated into biochemical routes that connect sugar and other input nutrients to the final desired target molecule (Routes layer), which introduces a vast combinatorial challenge. The fourth step (the Design layer) involves choosing which biochemical reactions to physically engineer within the strains. Next (in the Proteins layer) the Automated Scientist software decides what specific proteins to introduce into the cell or modify (in the case of native genes). Finally, it adds regulation and genomic context to the design (Layouts layer) and fully specifies the genotype down to the base pair level. Details on each of these layers are described in the remainder of this paper.

**Figure 3.**
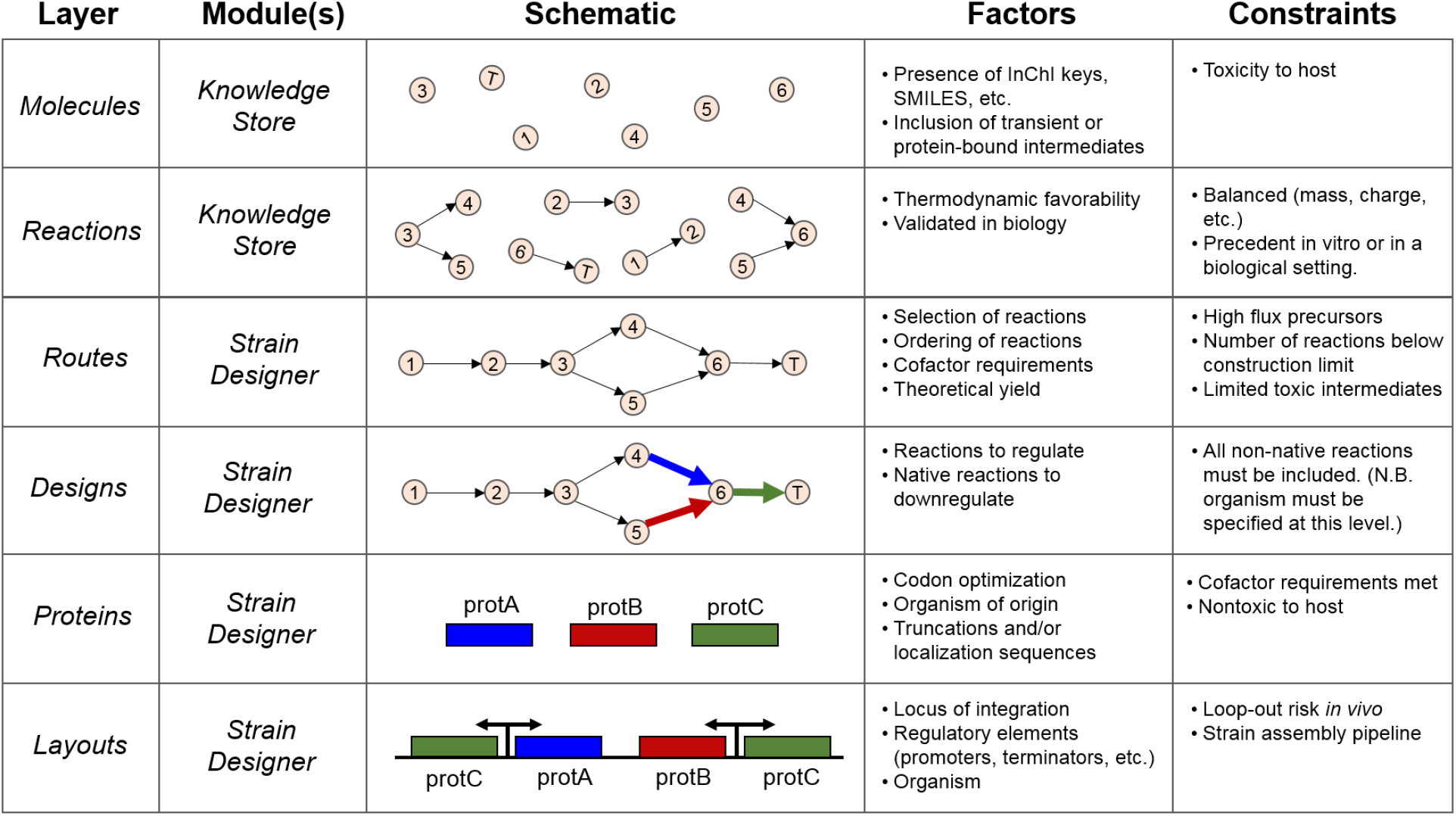
The Lila platform makes decisions at multiple distinct levels, from the biomolecules and reactions to consider to the final genomic layout. Each of these levels has a number of factors that can be experimentally modified as well as a number of constraints that must be considered.

### Lila’s Algorithm Generated Routes to 454 Diverse Biochemical Targets

To establish what biochemical reactions are permissible within the chosen microbial host, we populated the Knowledge Store with over 20,000 biochemical reactions, including those from the MetaNetX database^36^, into a repository we refer to as the Amyris Universal Set of Reactions (Figure 4A). We next annotated these reactions using a combination of public data and in-house curation, allowing Lila to filter challenging reactions based on their thermodynamic infeasibility, lack of experimental support in the literature, or other factors. Lila’s Route Finder algorithm then stitched these reactions together into a biochemical route from glucose to the target molecule of interest (Figure 4B) and optimal routes for 454 molecules were identified for each model organism (Figure 4C).

**Figure 4.**
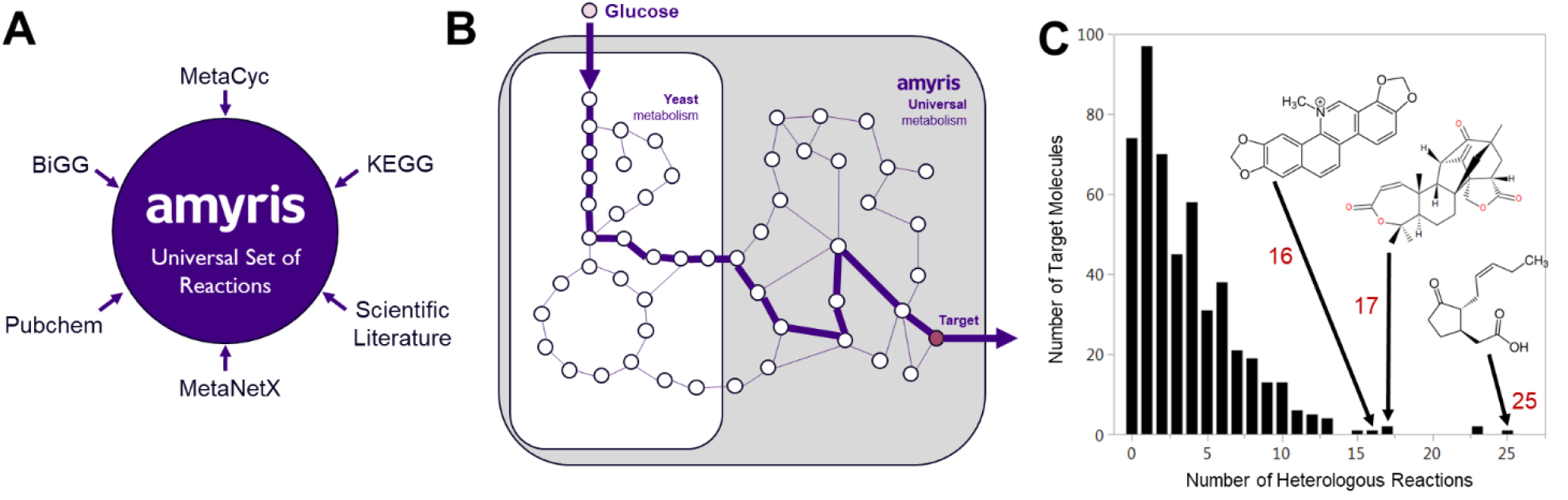
Automated route finding resulted in biochemical pathways going from glucose to 454 target molecules in both S. cerevisiae and E. coli. (A) Combing reactions from a variety of public databases and in house efforts resulted in a universal set of over 2.9 million compartmentalized biochemical reactions. (B) Biochemical pathways derived from this set of reactions span both native yeast metabolism (embedded white box) as well as non-native reactions (grey region). (C) The majority of pathways require 4 or fewer non-native reactions to complete the pathway, but some molecules needed 15 or more heterologous steps.

For each target molecule, Lila’s Route Finder emits an ordered list of native and heterologous enzymes that catalyze successive reactions to obtain the target molecule. Most target molecules require 1-5 heterologous enzymes to catalyze the pathway (Figure 4C). For each enzyme represented as an Enzyme Commission number^37^ (EC#), however, there are dozens to hundreds of enzymes encoded by various organisms that all catalyze the desired reaction. To decide which enzyme to use from which organism, we devised an algorithm that retrieves all possible amino acid sequences from Uniprot^38^ corresponding to each EC#, and adaptively chooses *n* or fewer enzyme variants per EC#, where *n* is decided by an internal pipeline capacity model. Because the human procedure for selecting enzymes involves reading the literature, we implemented literature-mining algorithms that search for probabilistically enriched co-occurrences of any enzyme aliases and EC#s in all PubMed abstracts, and detect keywords such as organism names, whether the enzyme has been purified, expressed, or assayed (SI Figure 7). In addition, we constructed phylogenetic trees for each group of enzyme homologs (data not shown) and retrieved information from the BRENDA database^39^ for any published information about kinetic parameters such as *k*_*m*_ and *k*_*cat*_.

To design an algorithm capable of working on par with a human scientist, we interviewed a panel of Amyris scientists with a cumulative 100+ years of strain engineering experience. The questions in the interview probed what criteria they used to select pathway enzymes for testing, how they thought about codon optimization, how they identified peptide targeting sequences, etc. Table 2 of the Supplementary Information summarizes their approaches and provided a starting point for prioritizing which approaches to automate in code. In a period of 12 months, this algorithm selected 1,850 enzymes for gene synthesis, and annotated them for correct insertion into hundreds of strains making molecules of interest.

**Table 1.**
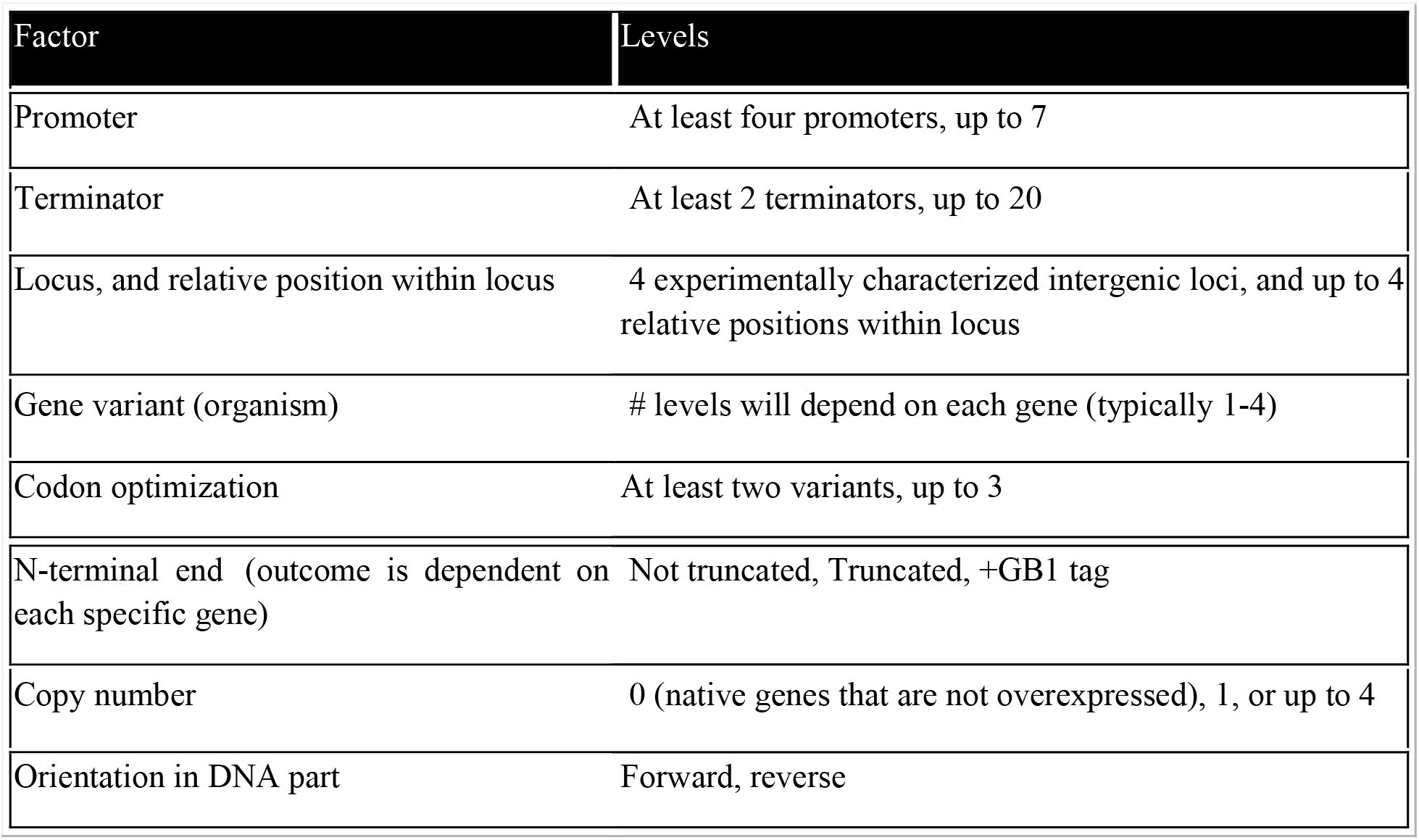
Factors and levels used in combinatorial strain design.

**Table 2.**
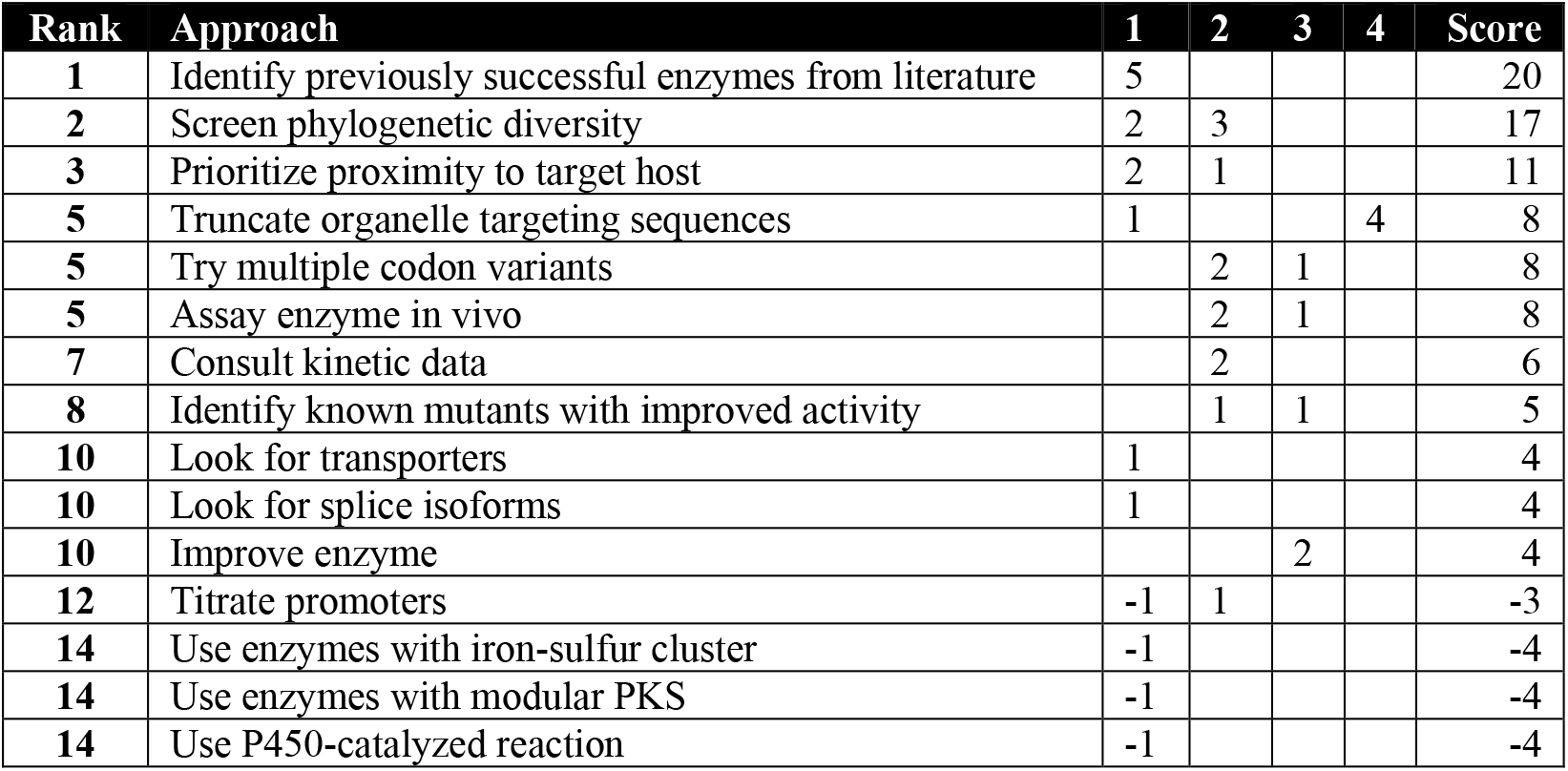
Ranked-choice scoring of number of strain engineers who would take each of the enzyme selection approaches as their first, second, third, or fourth choice. A -1 indicates that it is a strategy that they would explicitly avoid at first/second/third/etc.

**Table 3.**
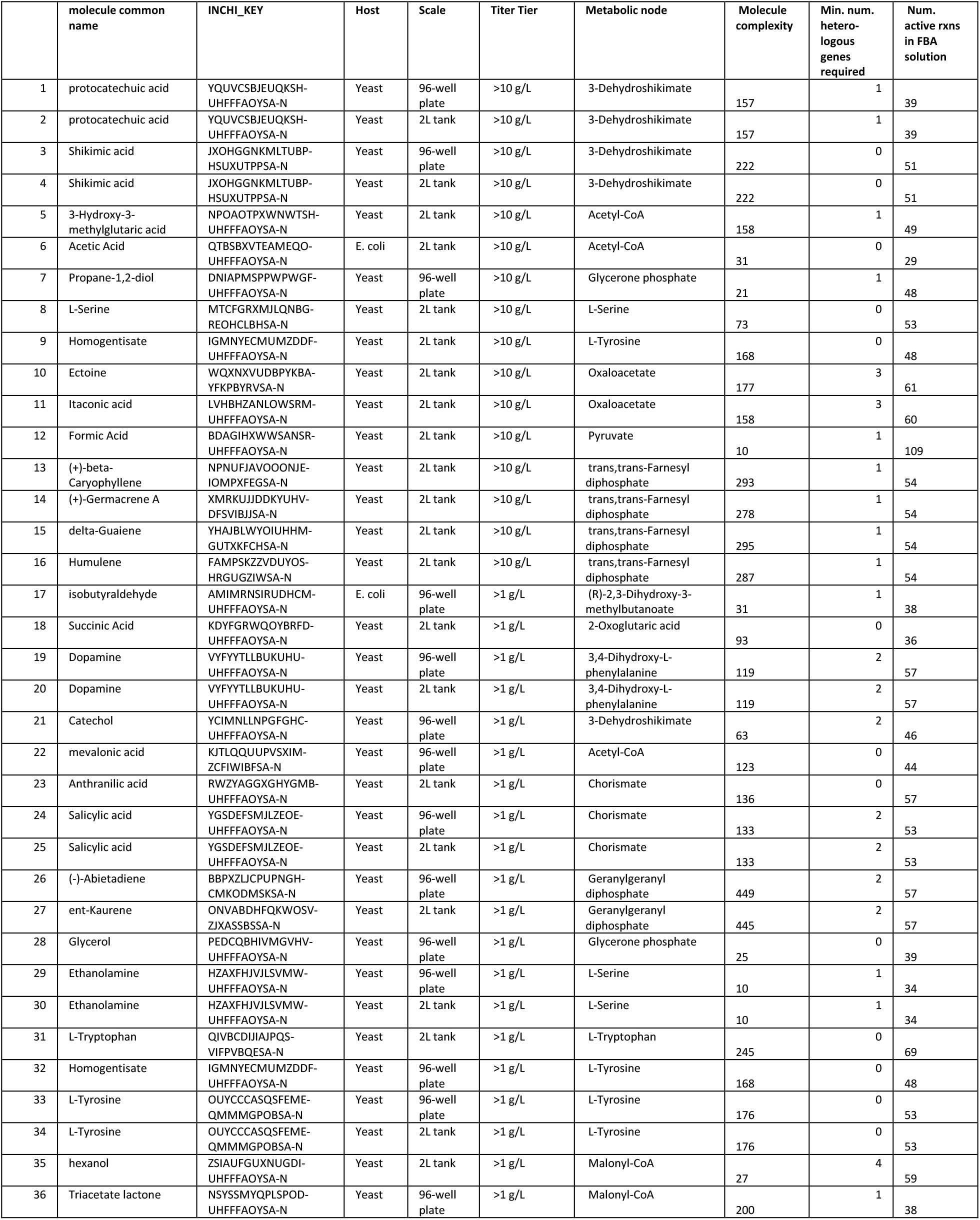
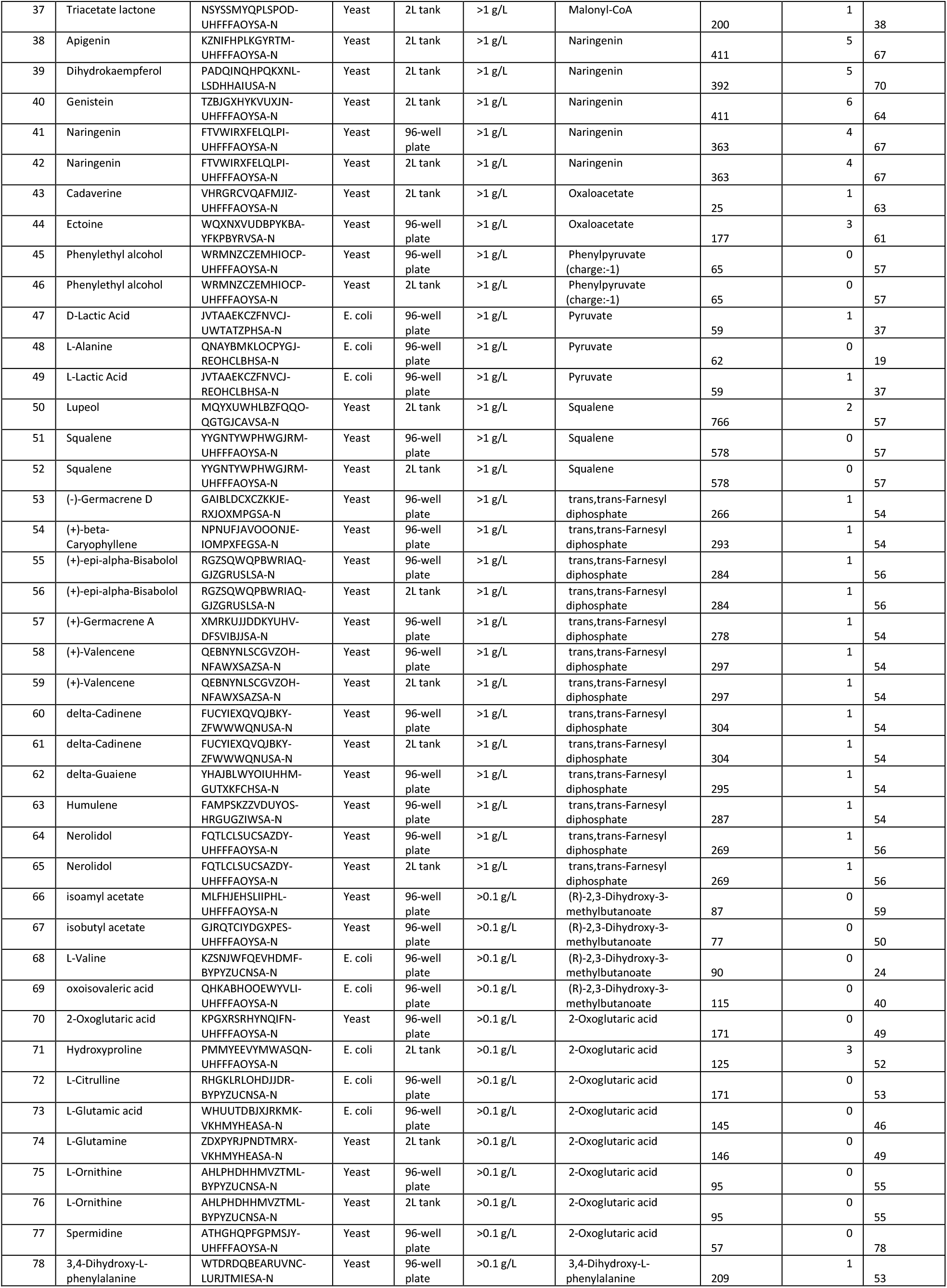
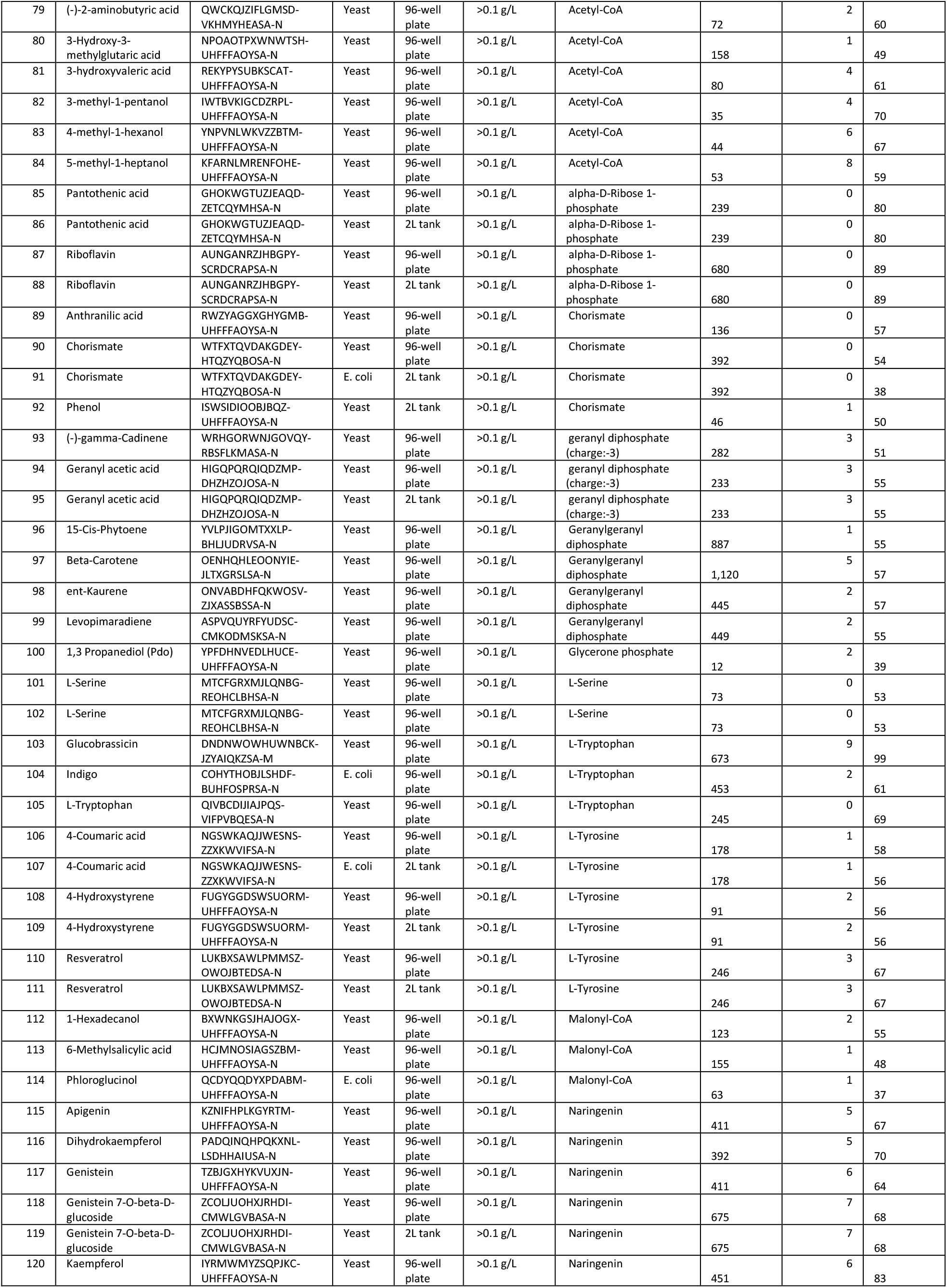
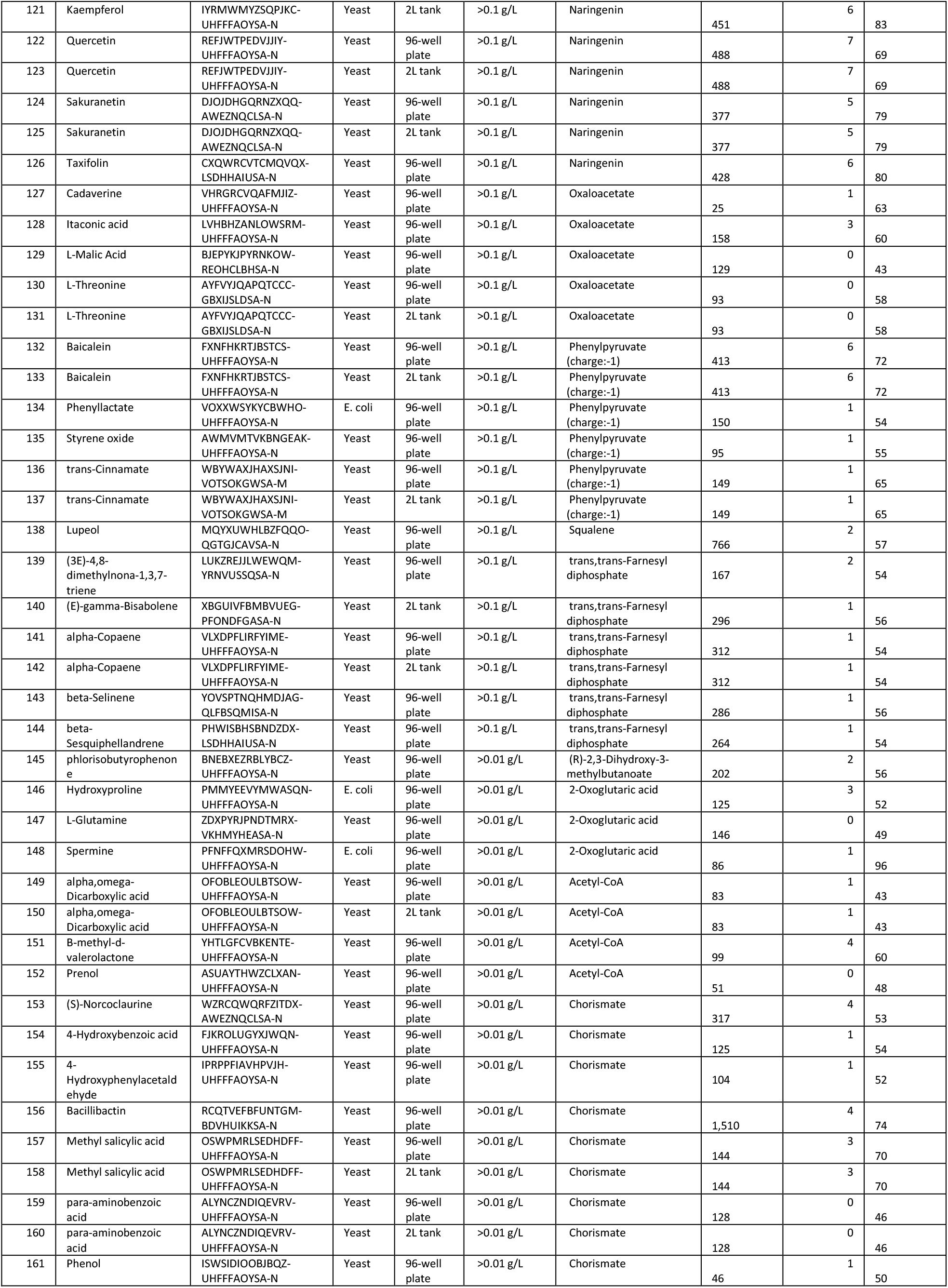
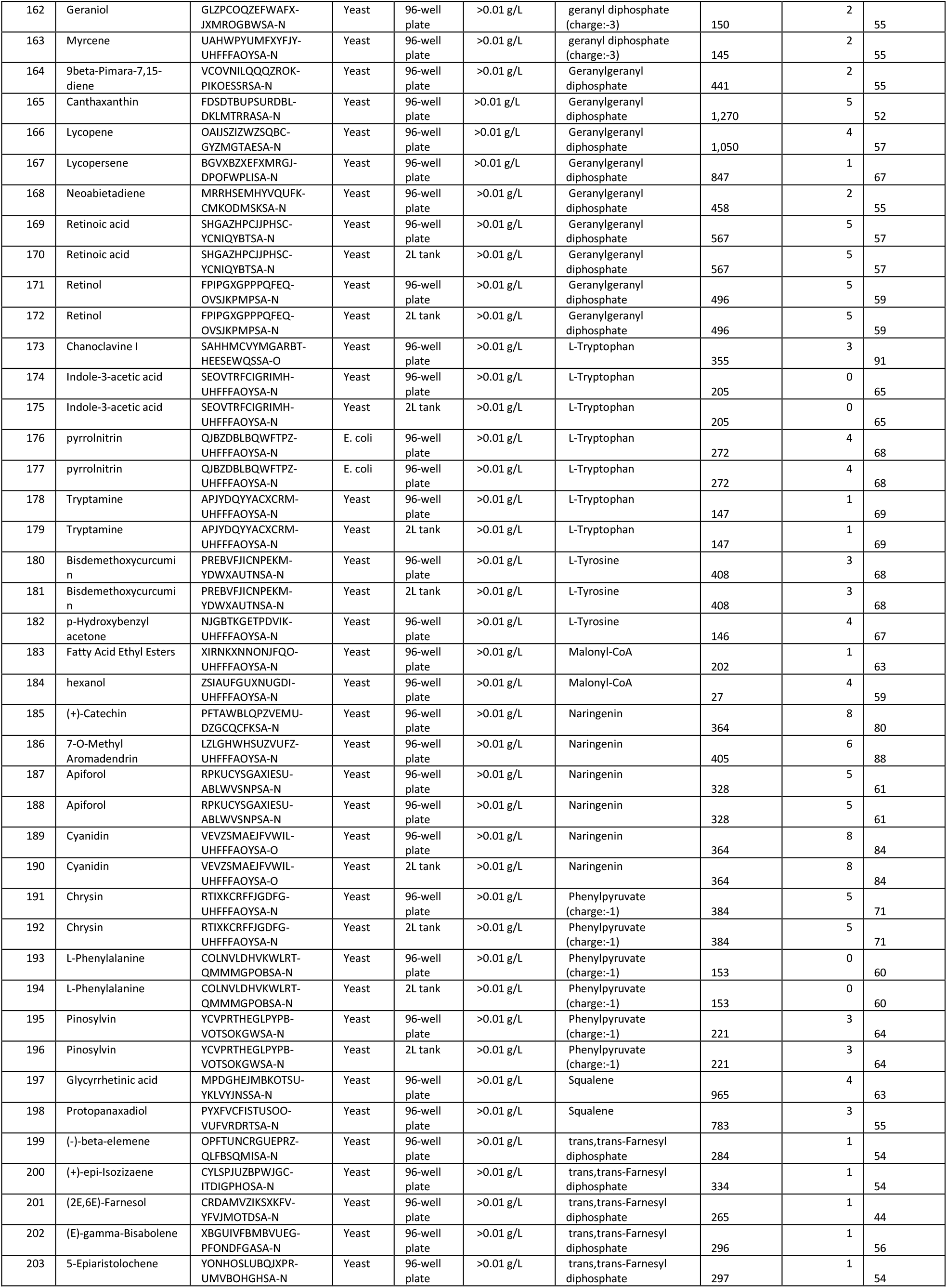
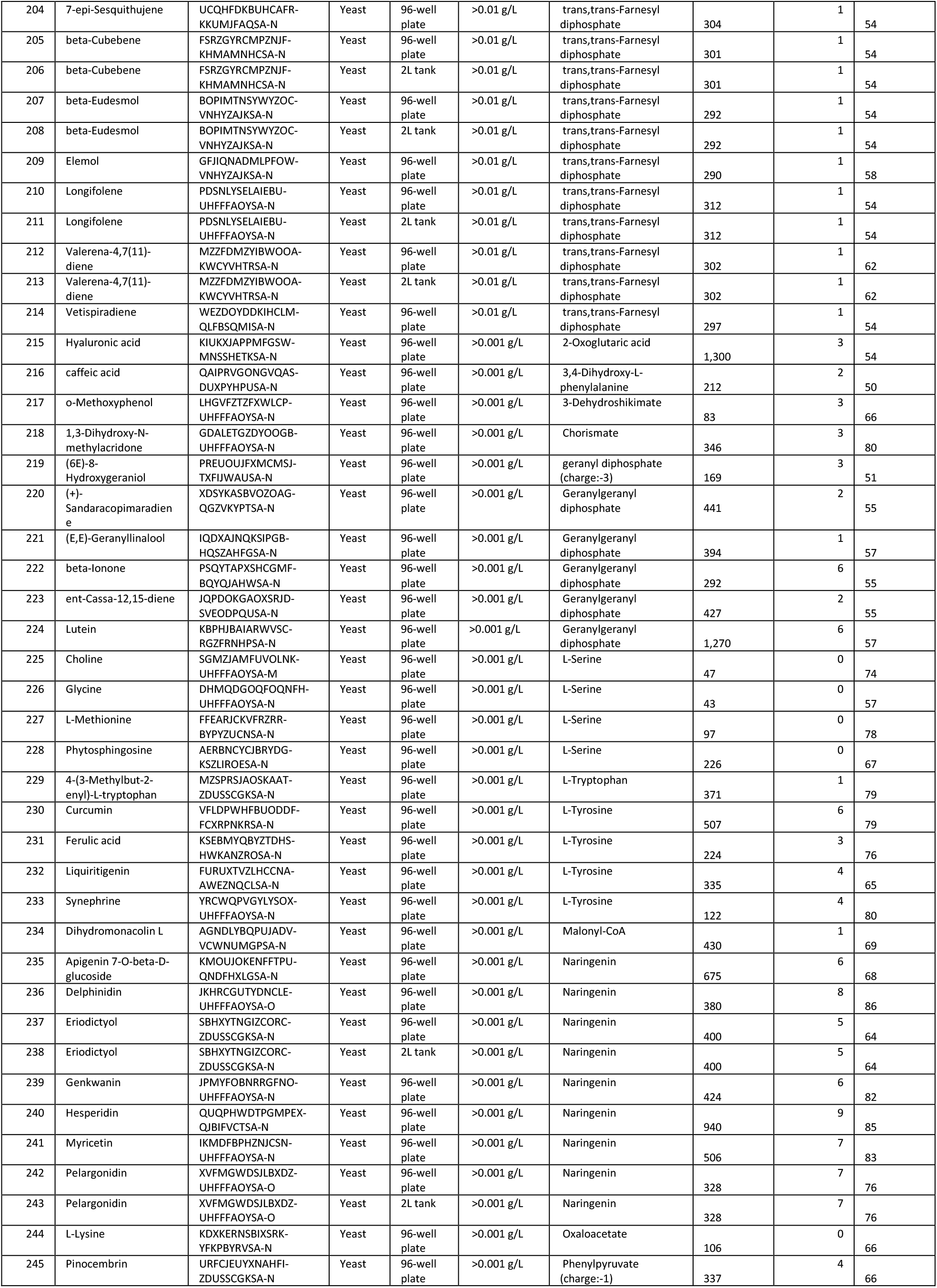
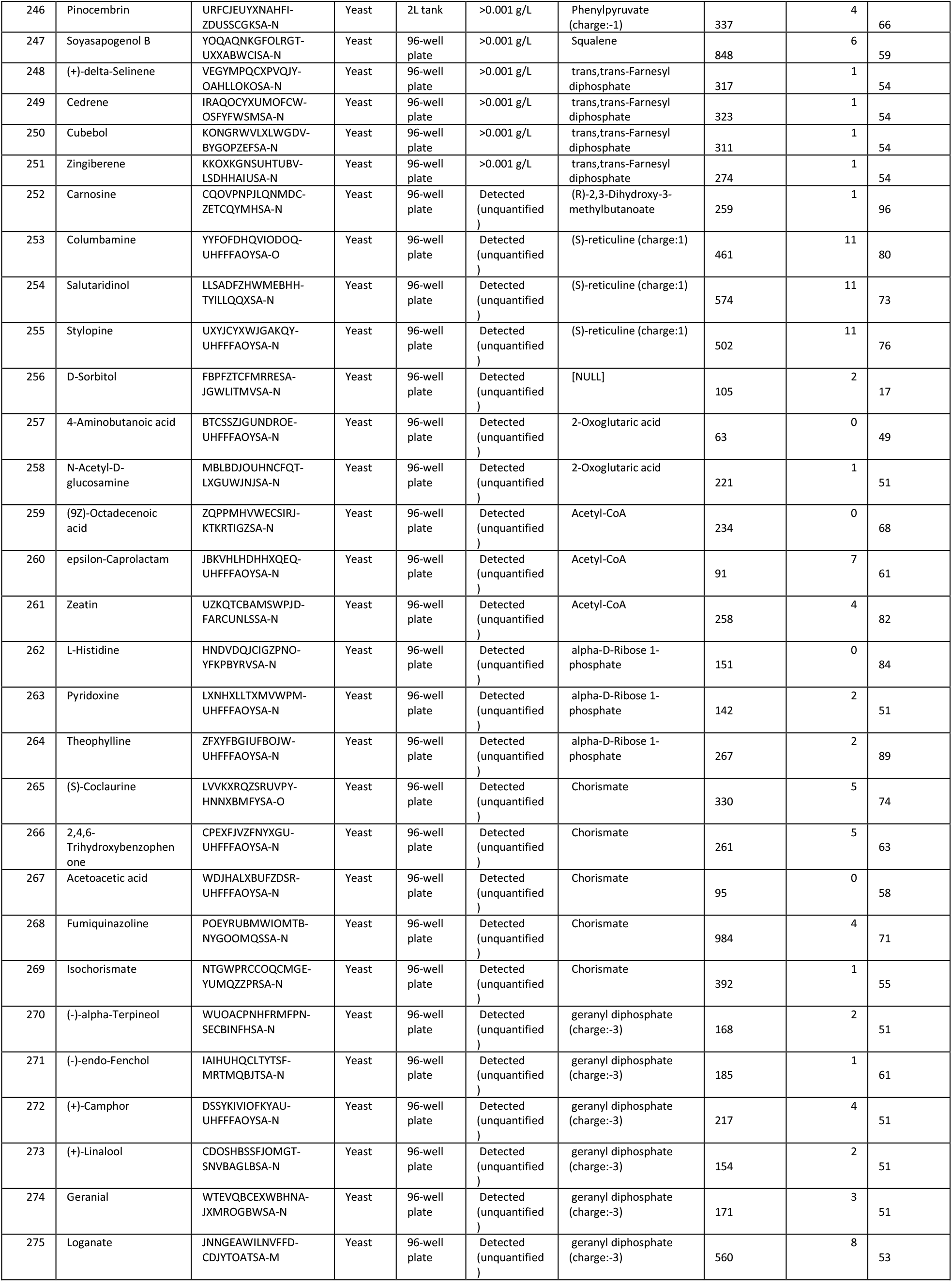
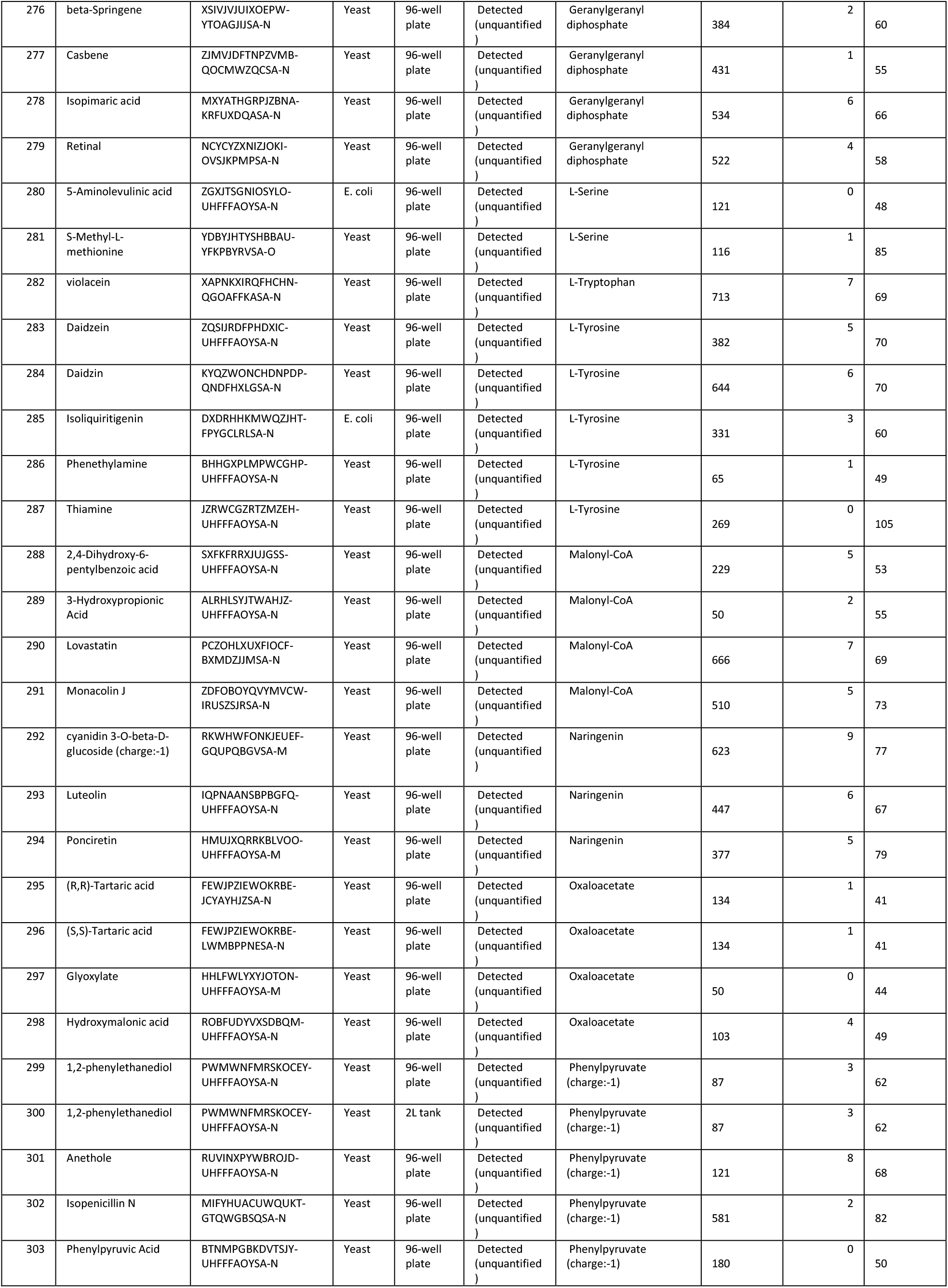
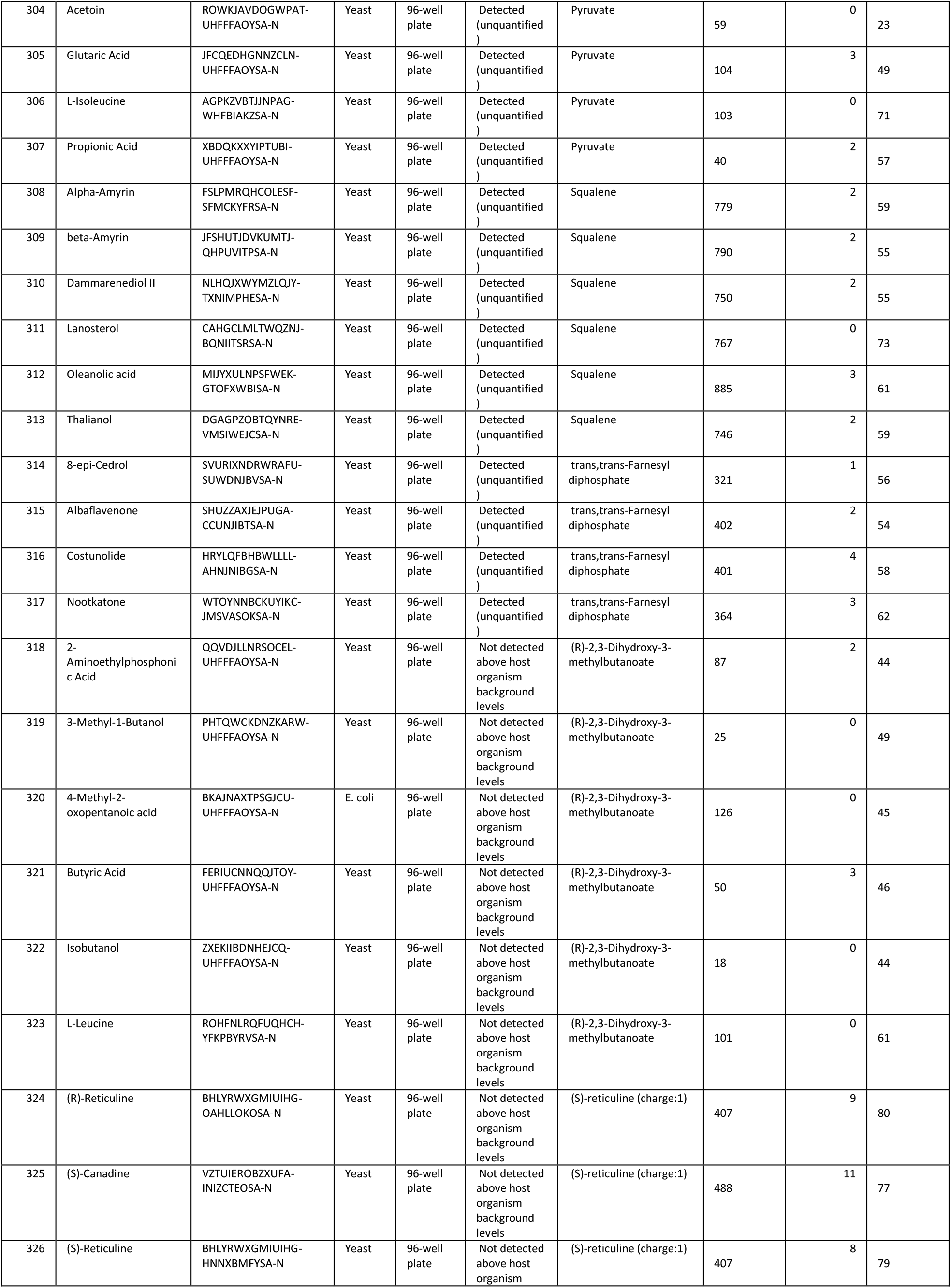
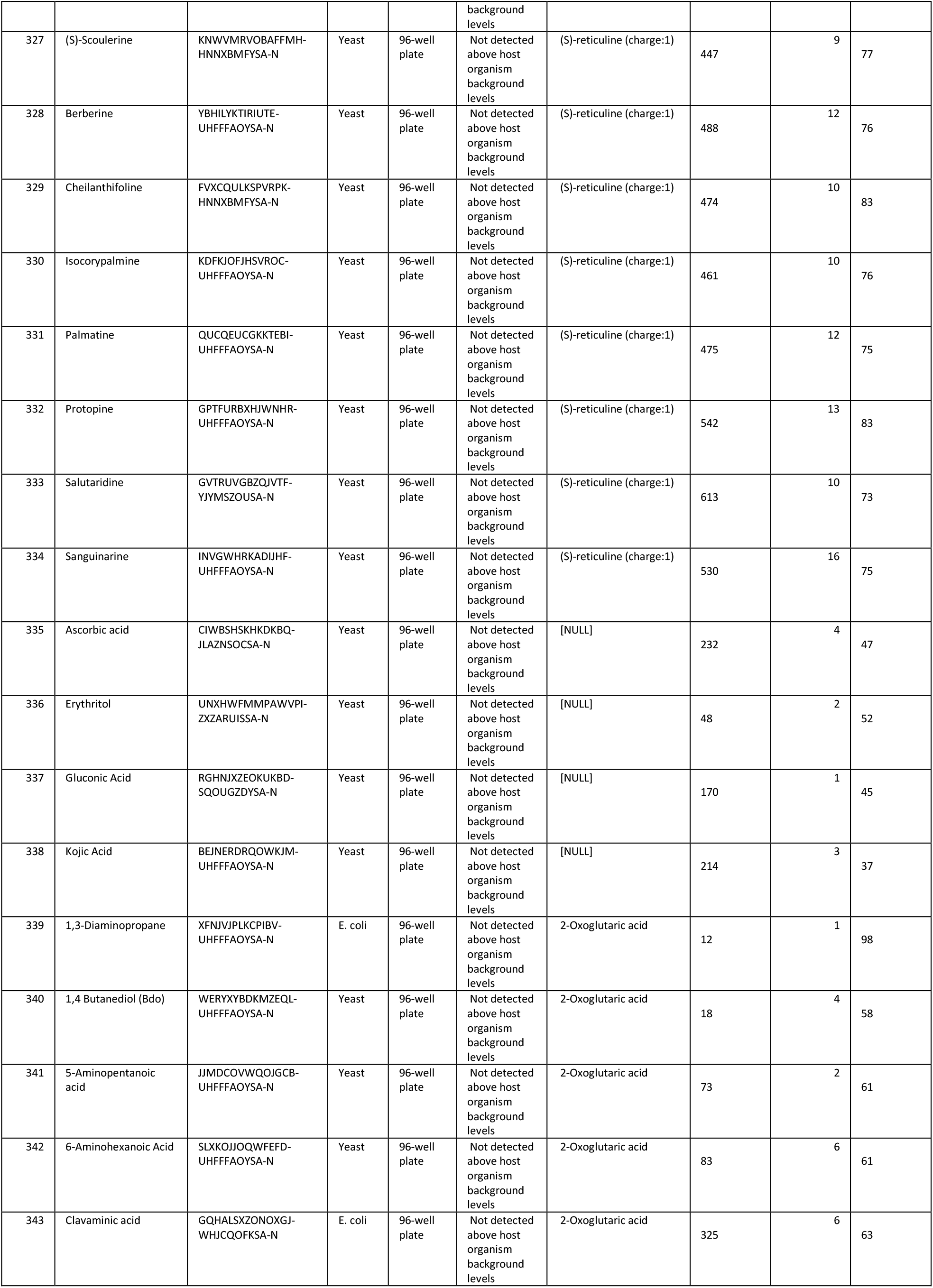
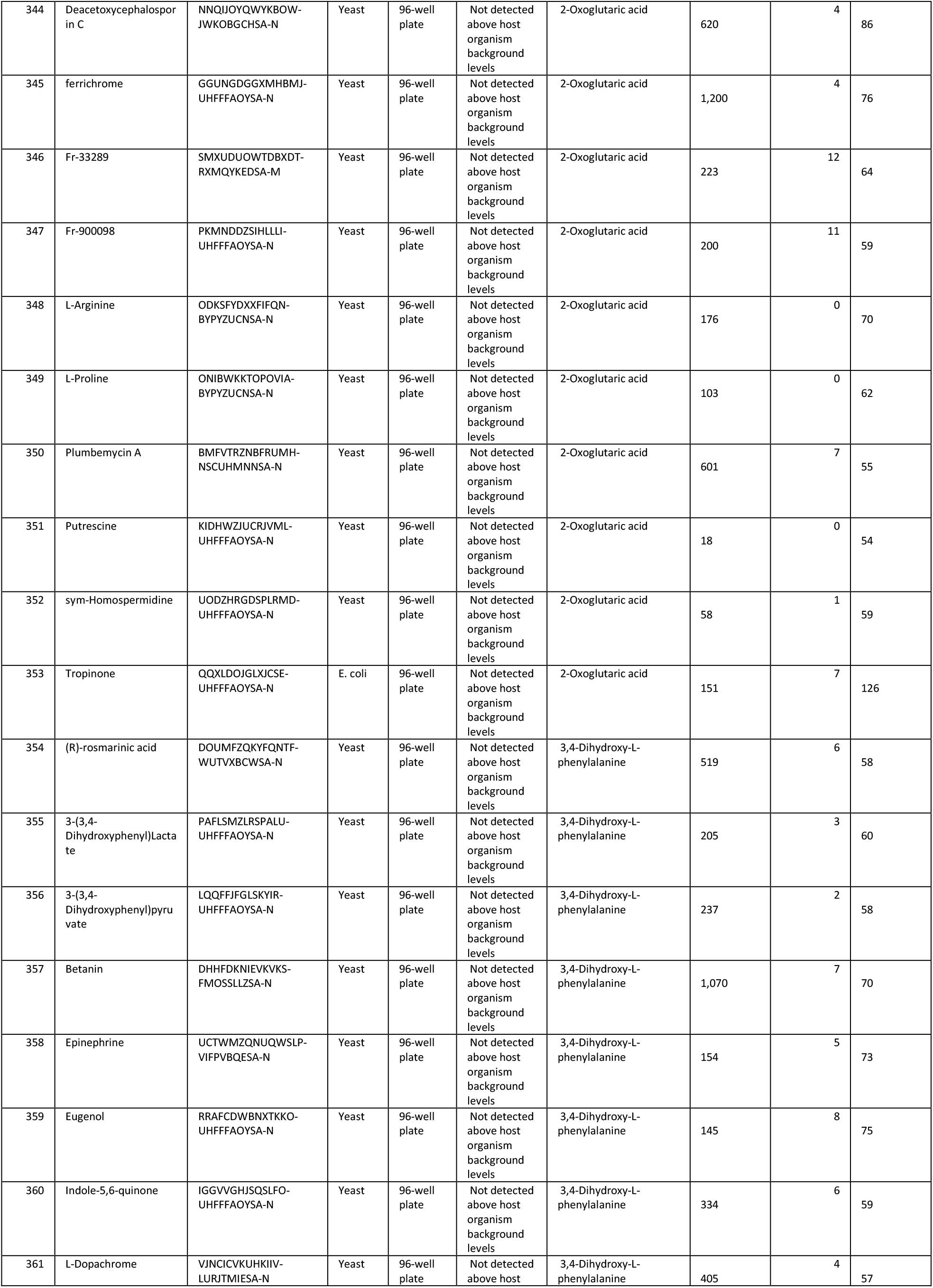
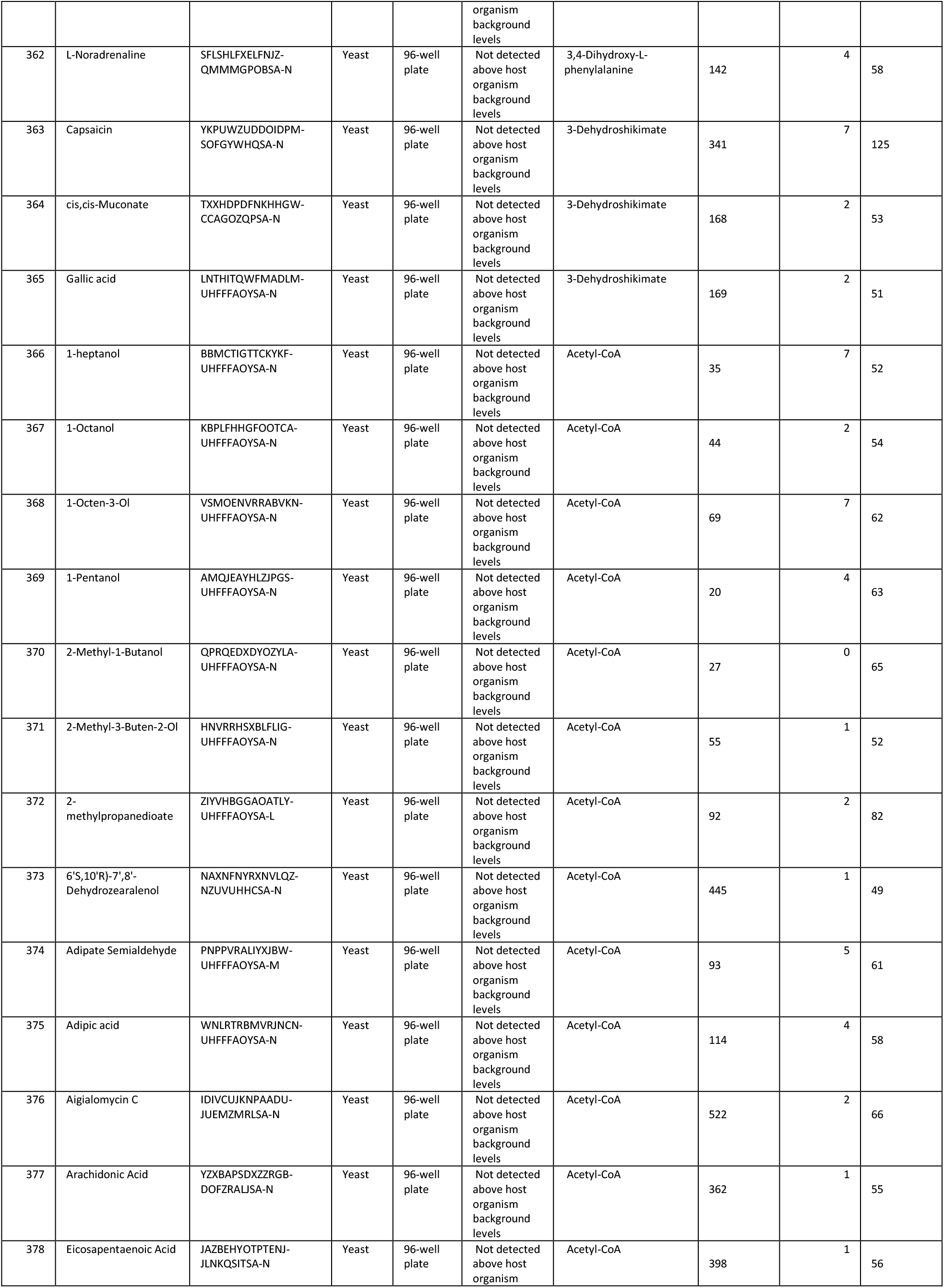
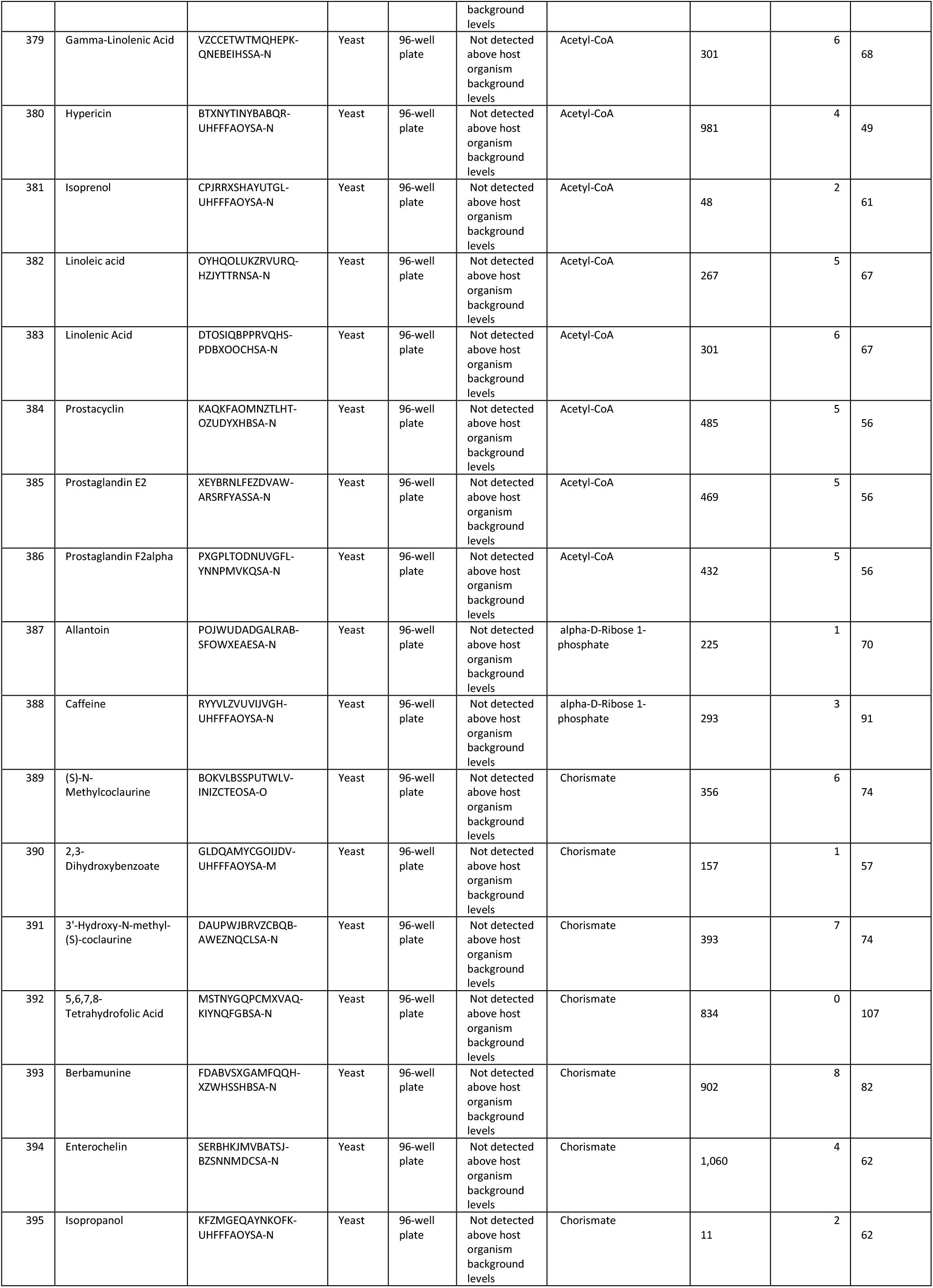
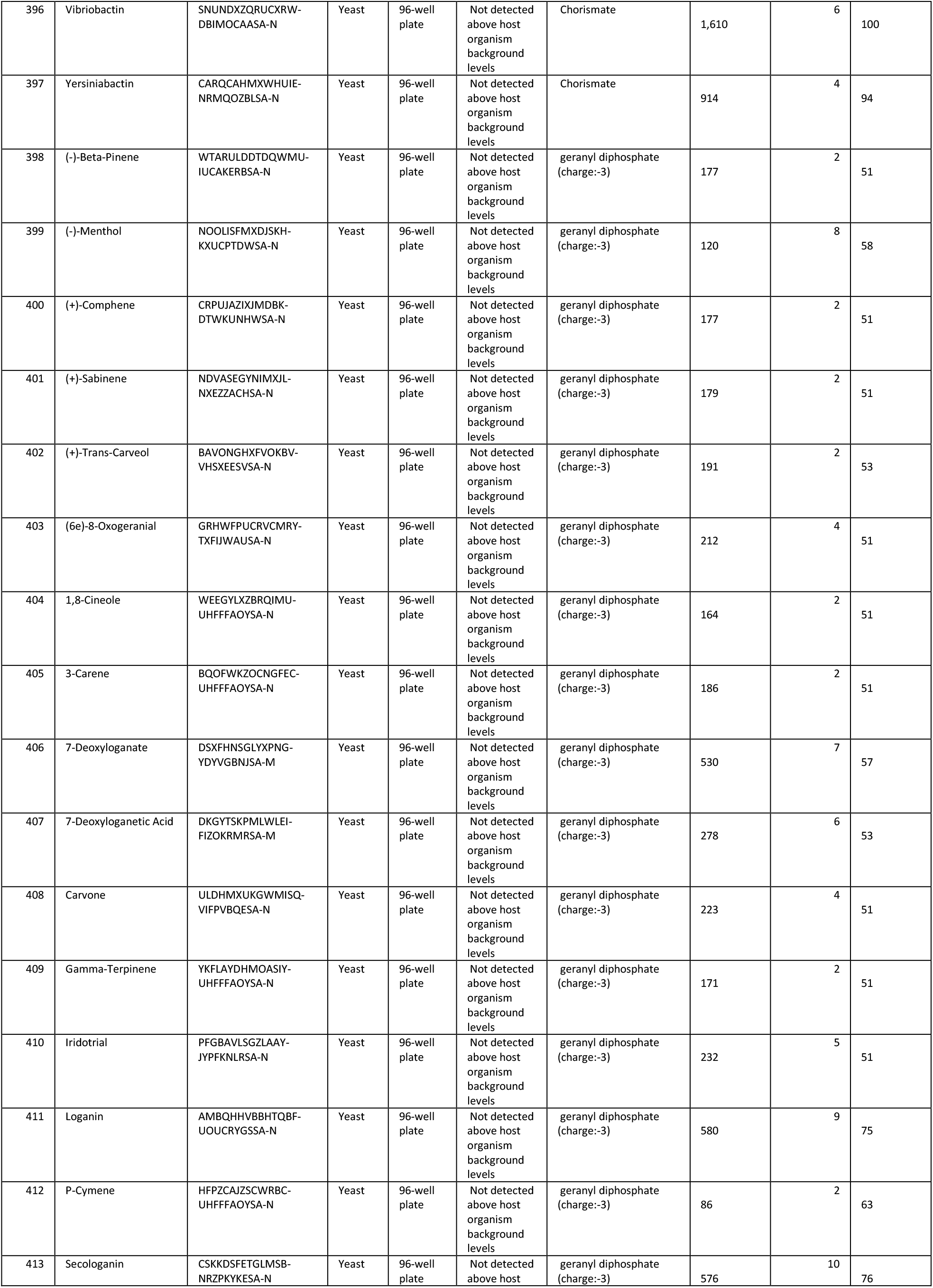
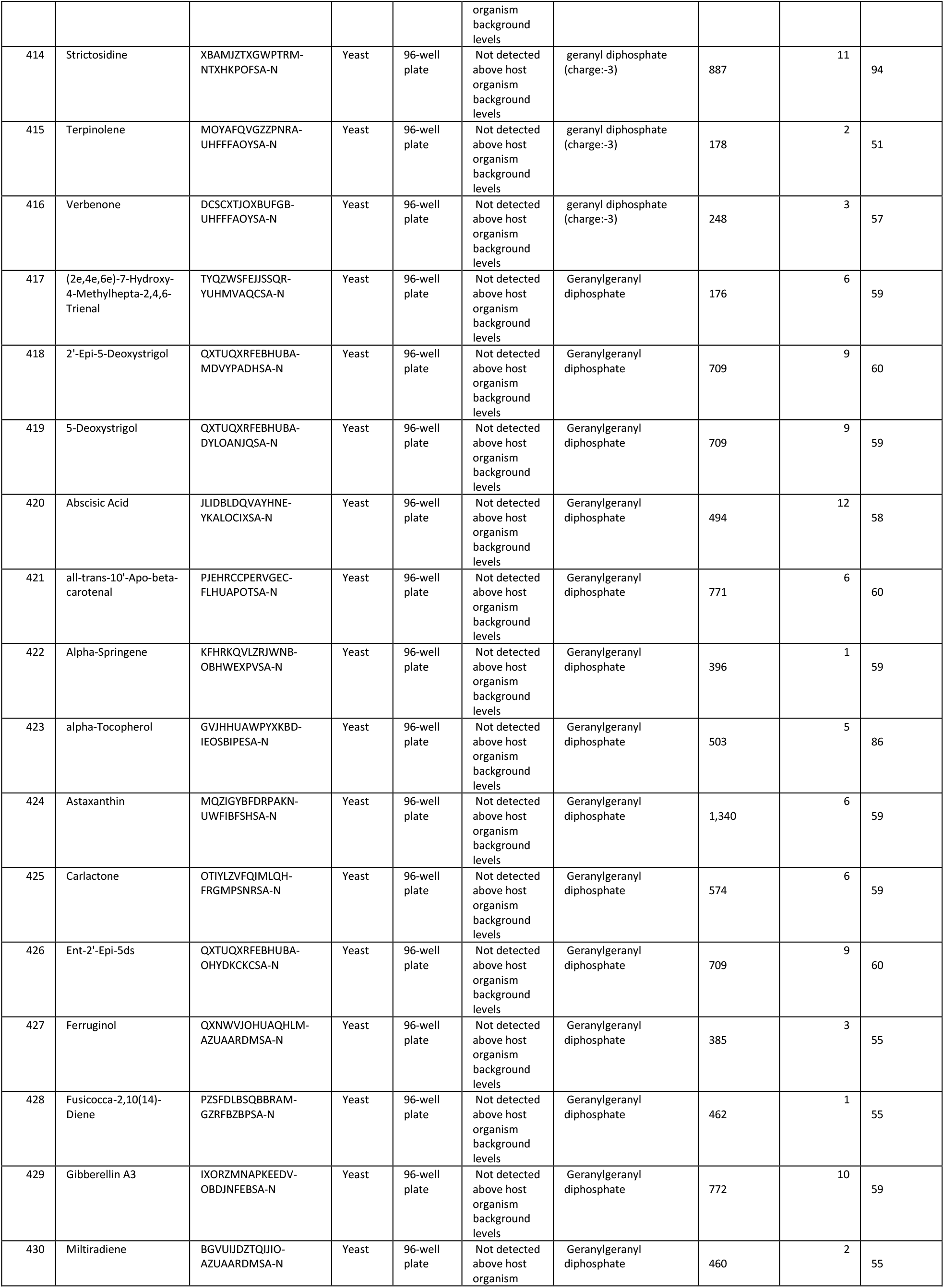
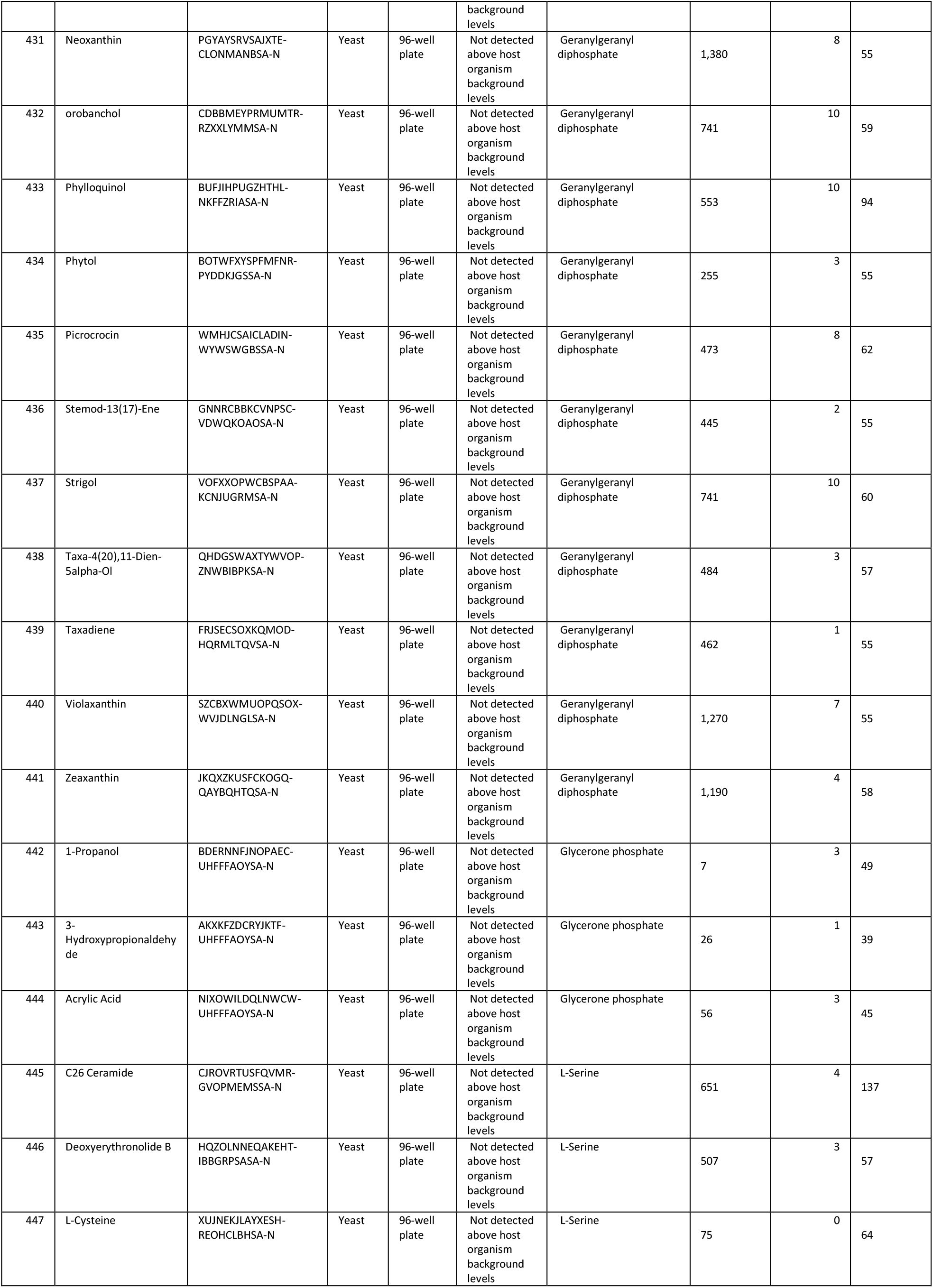
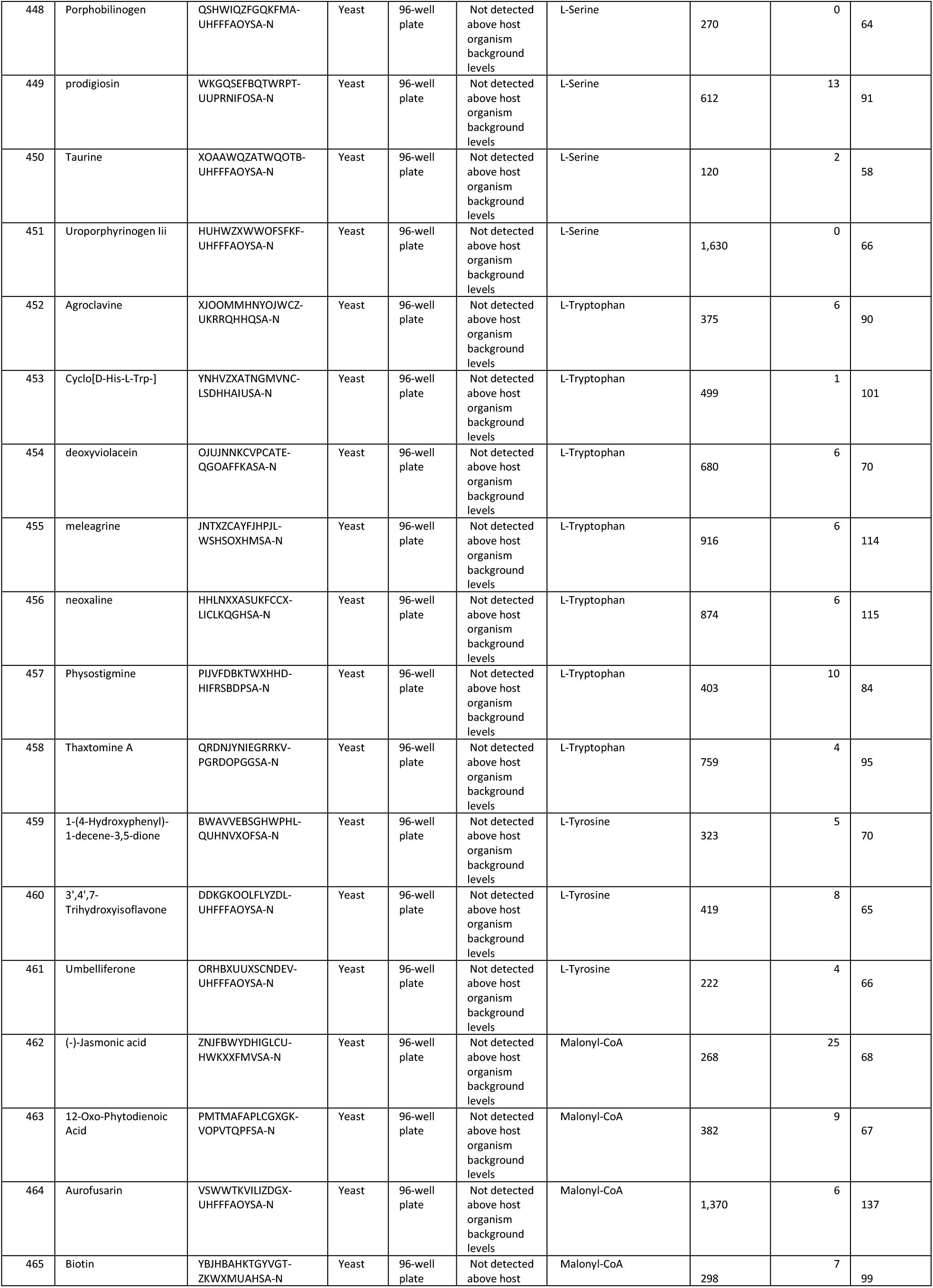
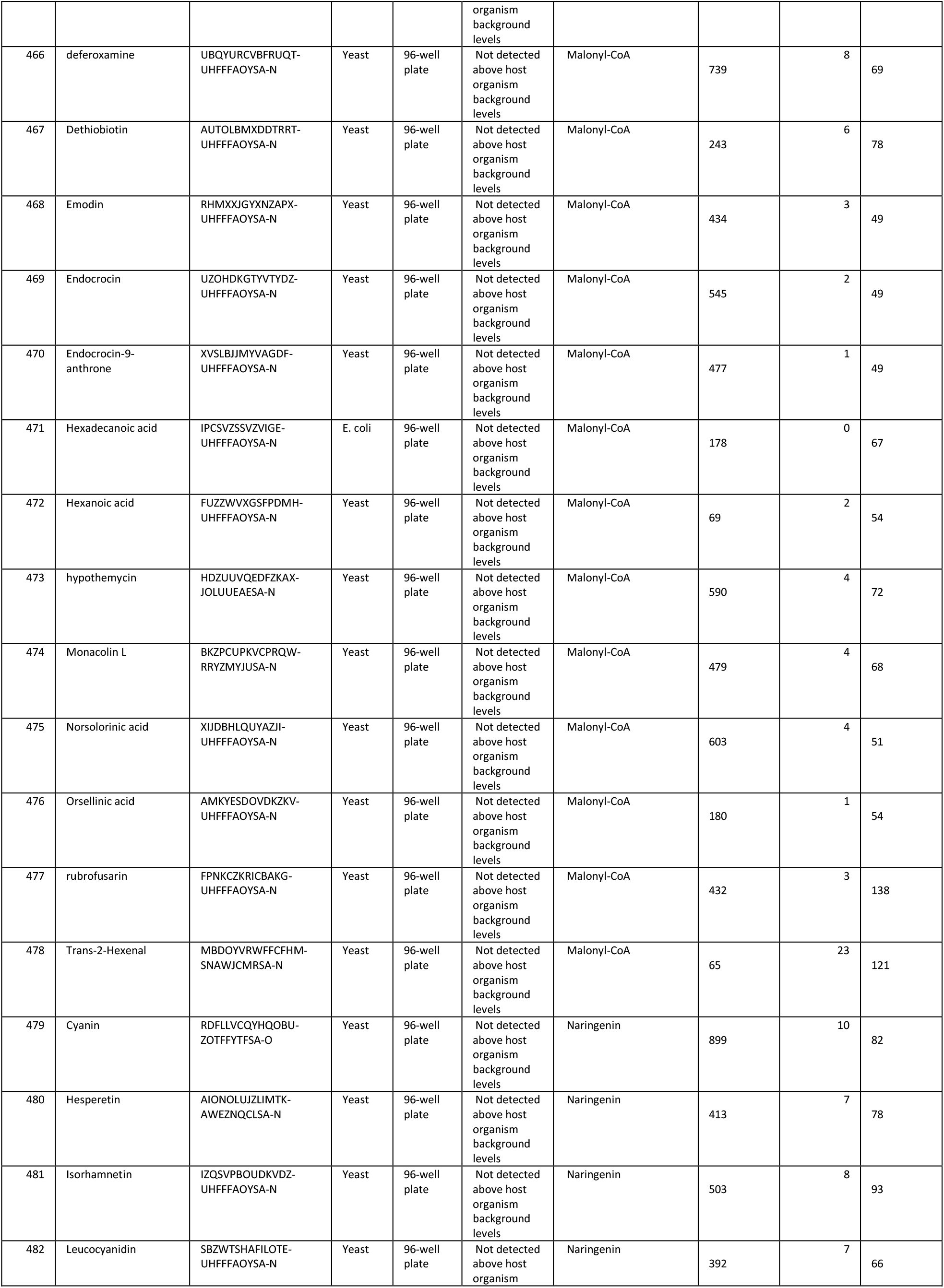
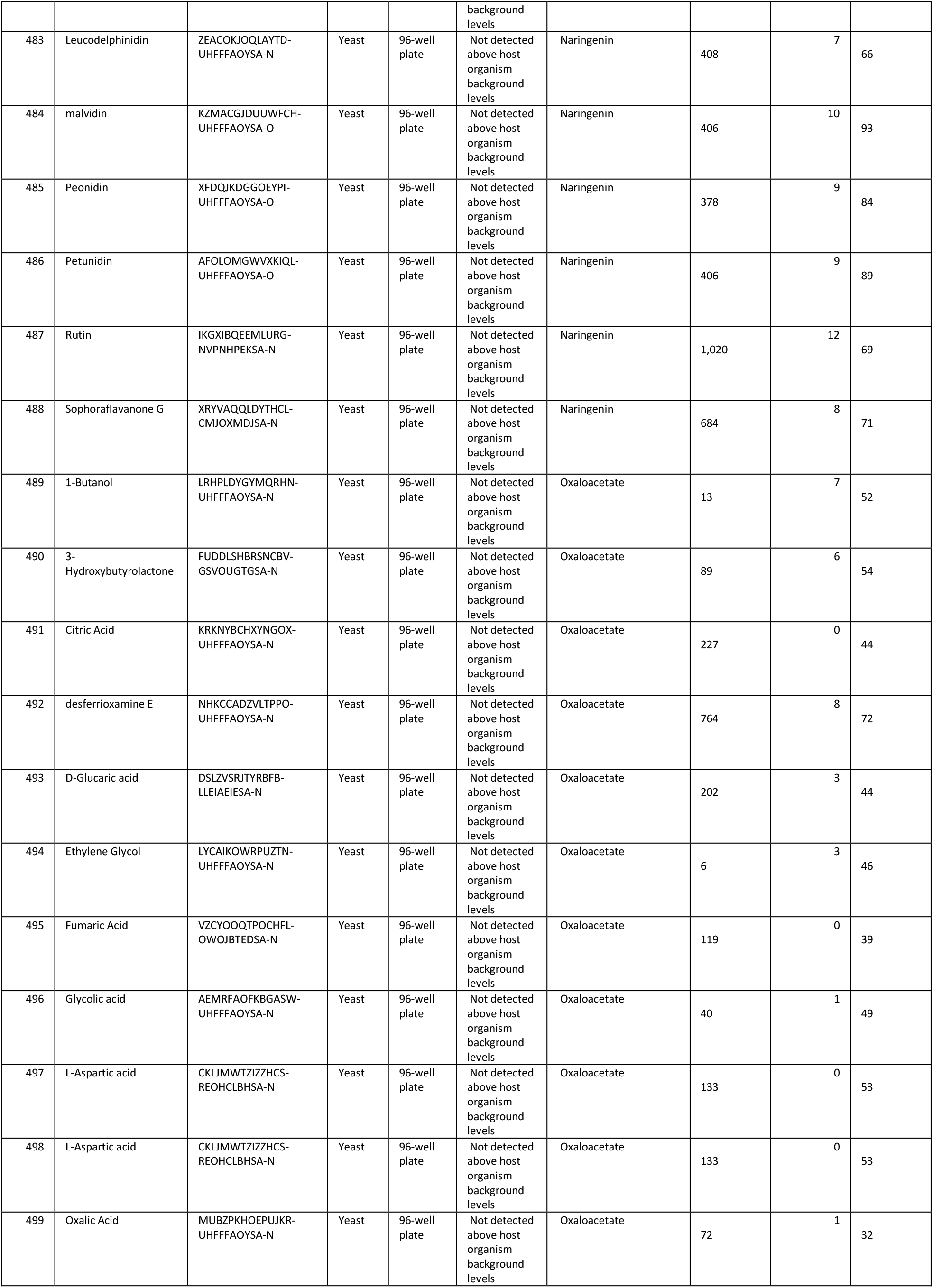
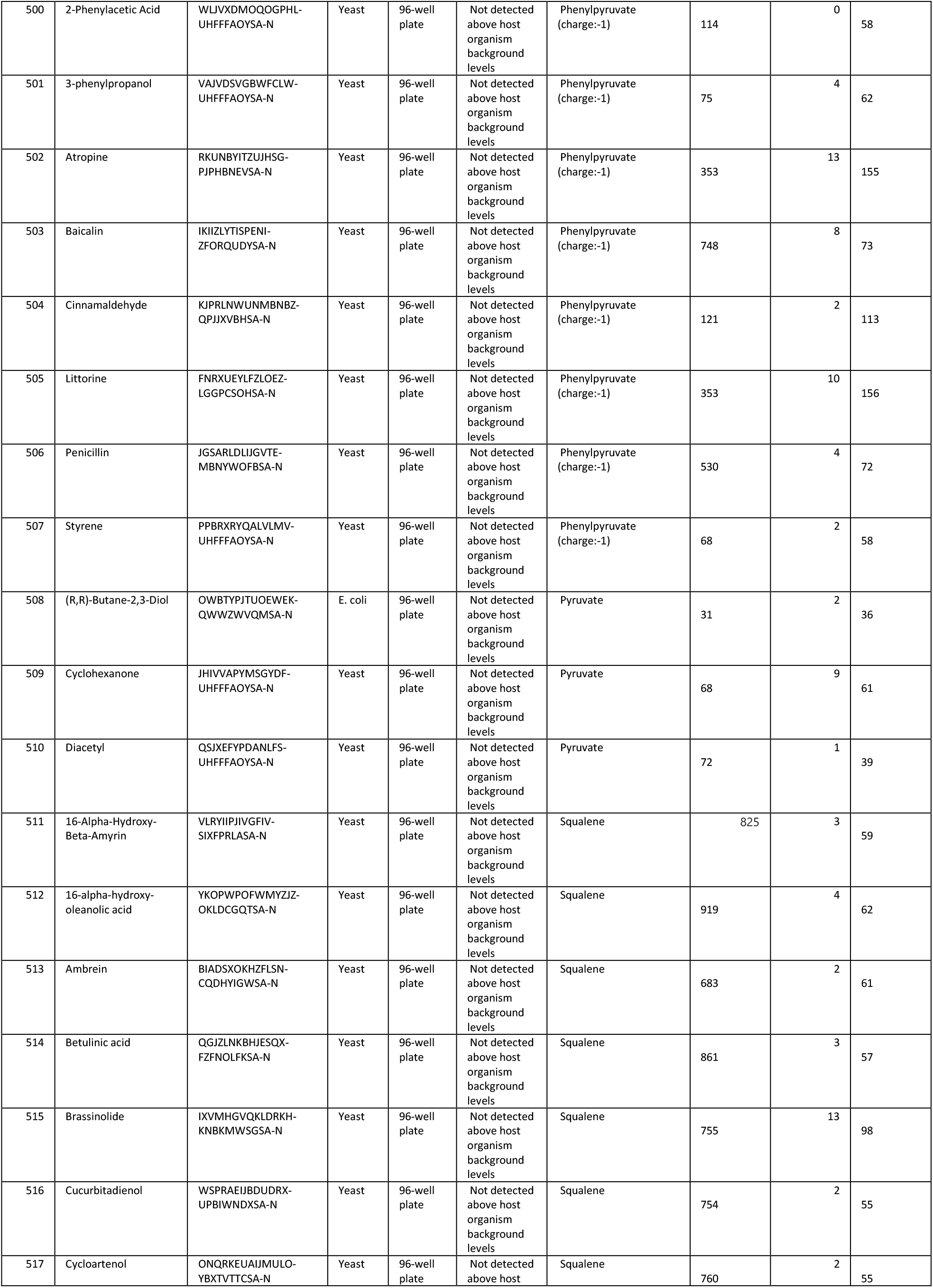
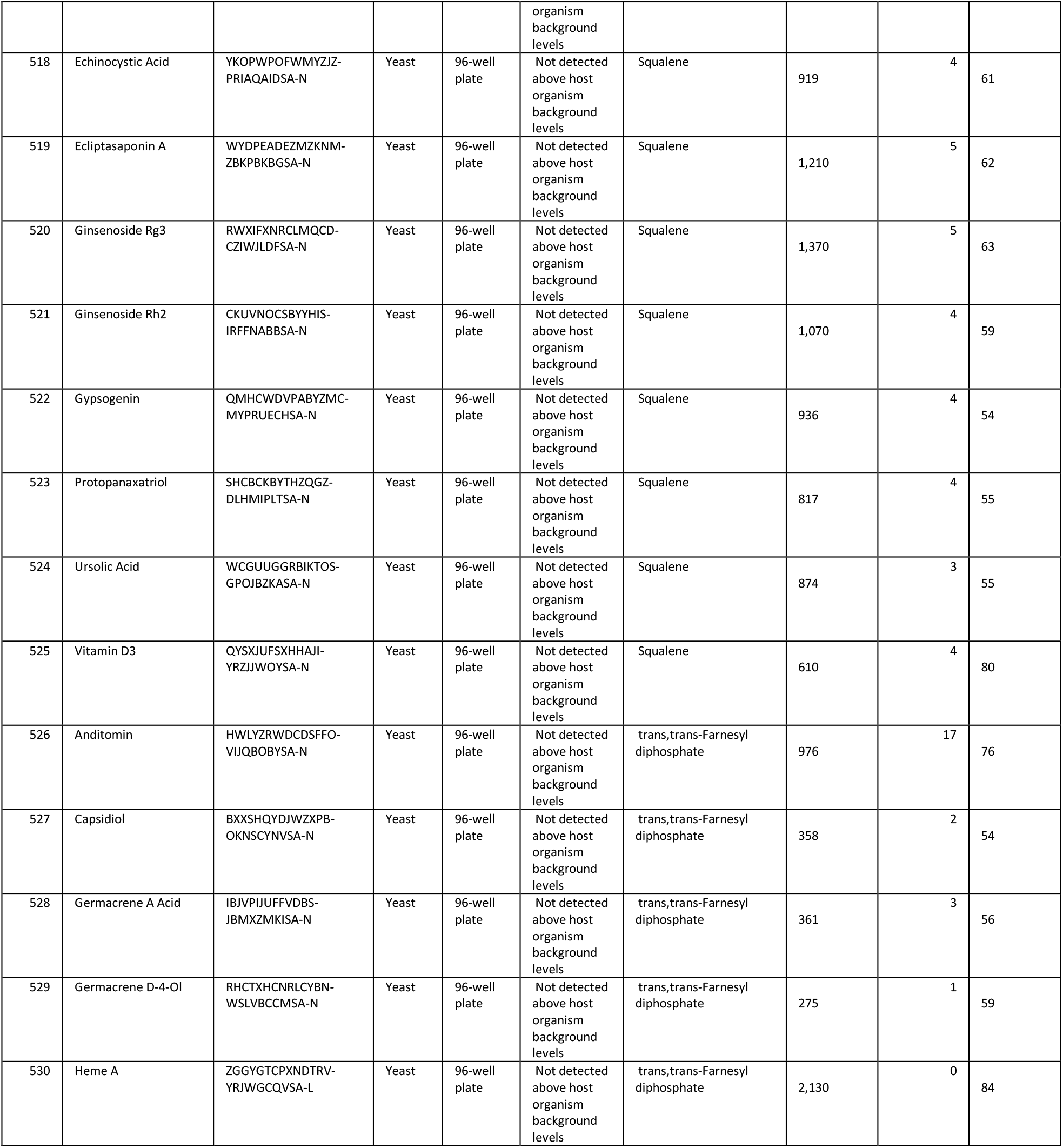
List of all molecules attempted, the highest titer tier observed, the organism the highest titer was observed in, whether the titer was observed in a plate of fermentation tank, the molecule complexity, the number of heterologous genes required to produce the compound, and the overall number of active reactions required to achieve maximal yield in the FBA solution predicted by Lila’s Route Finder module.

### Lila’s Strain Designer Explores Combinatorial Genetic Space Using Design-of-Experiments and Linear Optimization

Following the identification of a biochemical pathway and associated proteins, Lila’s next challenge was to convert these high-level choices into genomic layouts (Figure 3, last row). This step involves making decisions about genomic location (e.g., what location in which to integrate each selected protein), regulation (e.g., which promoters and terminators to include), and engineering feasibility (e.g., whether tandem or inverted repeats will be allowed, to ensure genetic stability).

To explore this vast combinatorial design space we employed a Design-of-Experiments based approach. As there are many flavors of designed experiments^40^, we generated a short-list and filtered it to variants that could be practically constructed by Amyris’ strain engineering pipeline and could plausibly generate metabolic pathway variants, which generally have categorical or constrained factors. The short-list includes *(i)* full factorial designs: *n*^*k*^ runs for *k* factors with *n* levels each, typically two levels, main effects only, *(ii)* fractional factorial designs: *n*^*k - p*^ runs, for *k* factors with *n* levels each, where *p* is the number of possible interactions between factors, *(iii)* definitive screening designs: ∼*2k* runs for k factors with up to three levels each, assessing main effects & interactions, and *(iv)* Box-Behnken designs: response surface methodology with up to three levels per factor, assessing interactions. The list of factors and levels are detailed in SI Table 1.

Finally, we tied this Design-of-Experiments framework into a genotype specification so that combinatorial designs could easily be specified without needing to spell out each design variant afresh in code for each experimental cycle. Generating genotypes requires translating the high-level designs, which largely consisted of a list of enzymes to insert, overexpress, down-regulate, or remove, into low-level DNA constructs and strain specifications for Amyris’ Automated Strain Engineering Pipeline (ASE). Communicating a design specification to ASE is done via Genotype Specification Language^41^ (GSL), but for the specific task of automating thousands of designs, we needed to encode repeated sets of complex GSL statements (higher-level design constructs) into simpler function calls. We therefore implemented GSL functions to express a given number of genes at a given locus, drive a given gene with a given promoter, and so on. In consultation with biologists on the design of the ratified base strains for each microbial species, we encoded several GSL functions in advance that could insert the heterologous genes into the pre-existing landing pads.

Because our pipeline has finite capacity to construct strains and we were pursuing many targets at once, our samplings of these combinatorial spaces were often quite sparse. As a result we frequently began with a small number of strain designs at diverse points throughout the combinatorial space, and iterated until we converged to fewer successful designs. Once an algorithm identified the proteins that should be overexpressed, we implemented a number of strategies to formalize the re-design^42^, trying different homologs of the protein, and finding a new route that avoids the protein altogether (if the protein cannot be further overexpressed or if no suitable homologs can be identified).

### Lila Generated Hit Strains Producing Hundreds of Non-Native Small Molecules that Span Yeast Metabolism

Once strains were constructed using the Strain Designer module and measured for product titer, it was possible to begin the strain optimization phase. To identify and correct sub-optimalities in a biosynthetic pathway within a live strain, several factors and corresponding measurements were considered. First, the amount of flux passing through each metabolic node or reaction in the wild-type strain guides the type and degree of genetic or process-level perturbation needed. The expression of pathway enzymes is also important, and one wants to avoid scenarios in which pathway enzymes have poor expression while competing steps have plentiful expression. Third, the pool size of a given metabolite matters: the higher the substrate concentration, the higher the thermodynamic driving force towards the target molecule of interest. Thermodynamics (e.g., reference *ΔG’*^*m*^) plays a role particularly if there is an off-pathway step that is undesirable, but thermodynamically very favorable. Finally, while build-ups of pathway intermediates may in some cases represent wasted carbon not going to the desired product, and therefore undesirable, they may also be necessary in some cases to drive thermodynamically unfavorable reactions within a given pathway.

Each of these issues requires a different treatment in terms of the diagnostic approach. To close the loop between data analysis and strain re-design, we wrote a dozen algorithms that take an inductive approach in which data, rather than rules supplied by human domain experts, drive the design effort. We also relied on biologists to examine the strain designs being emitted by Lila and suggest changes or improvements. Phenotypic data such as next-generation sequencing, optical densities, titers, and different flavors of proteomics and metabolomics are mapped by these algorithms to extensively curated metabolic models derived from published models for *S. cerevisiae*^43^ and *E. coli*^44^. Conceptually, this work introduces a form of ensemble learning whereby multiple weak learning algorithms (i.e., simple algorithms with substantial error rates) are run in parallel to yield an overall lower error rate.

The results of this pipeline are shown in Figure 5 for the target molecule naringenin, a key platform node. Strains designed by Lila (Figure 5B) produced naringenin in a microtiter plate-based model at a wide range of output titers, from zero to >1.5 g/L (Figure 5A). To our knowledge, this titer of naringenin exceeds the highest published titer in the literature^45–48^.

**Figure 5.**
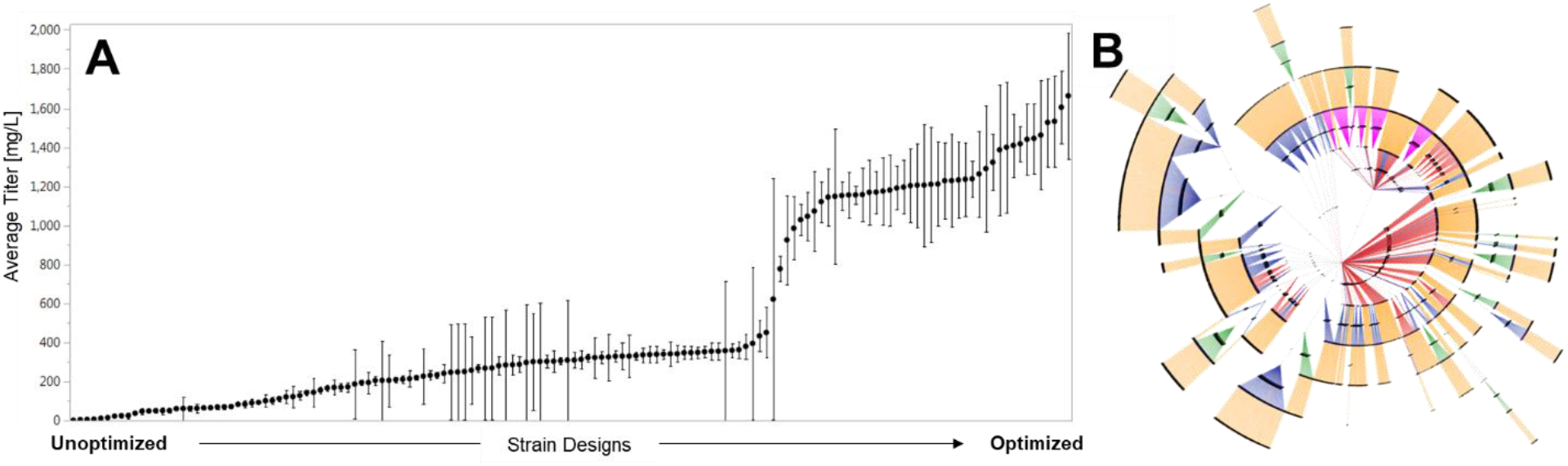
Successive strain and design improvements to produce the target molecule naringenin. (A) Titers in microtiter plates. Each dot represents the mean of all titer measurements for a given strain, and the error bars represent the standard error over N≥3 replicates. Only strains with N≥3 titer measurements are shown (n=163). (B) Comprehensive strain lineage map for naringenin strains (n=2,879) in a circular layout, with the common base strain in the center. Two nodes are linked by a directed edge color-coded based on the type of genetic modification used to modify the parent strain: promoter swap or knockout (green), integration of heterologous genes at one of three different genomic landing pad positions (blue, red, and pink), and plasmid curing (orange).

The overall results of tasking Lila to develop strains producing 454 different molecules are shown in Figure 6. Over the course of 105 partially overlapping DBTL cycles, 242 distinct molecules were detected at levels above the host organism’s native level, 140 of which achieved titers over 10 mg/L in microtiter plates, while 96 of which achieved titers over 100 mg/L. A subset of strains for 77 distinct molecules were tested in a 2L fed-batch bioreactors^49^ to further interrogate strain potential and generate material for applications testing. This often led to an order of magnitude increase in observed titer (Figure 6).

**Figure 6.**
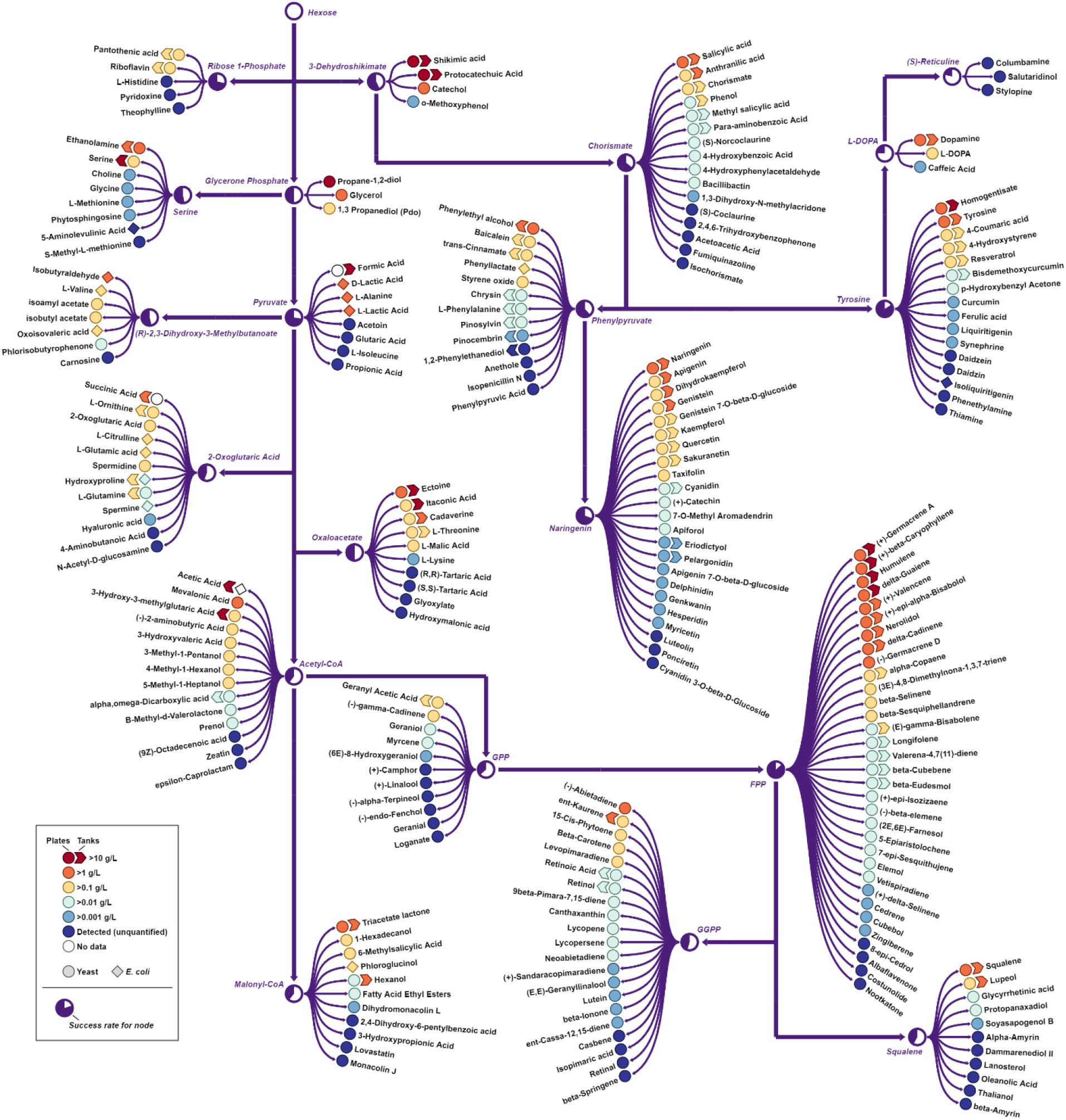
Titer results across metabolism show titer hits in 21 of 22 different metabolic nodes. A simplified metabolic map shows all targets with detectable titers. Success rates for metabolic nodes are the fraction of all targeted metabolites for which the molecule was detected in either 96-well microtiter plates containing 40 g/L sucrose^52^ or 2L fed-batch fermentation tanks^49^. Molecule titers were quantified by generic GC-MS or LC-MS/MS methods^51,52^. The map was generated using the Escher^55^ software package.

A comprehensive description of all attempted molecules, their relative molecular complexity^50,51^, and two measures of pathway complexity, can be found in the SI Table 3. We observe that more complex molecules and pathways generally resulted in strains with lower titers (SI Figure 8, Figure 9, and Figure 10). For example, the average molecule complexity at titer tiers above the >0.1 g/L threshold were all significantly lower than the average molecule complexity for molecules that were not produced at levels above the host organism’s native levels. However, for detection of any production that was above the host organism’s native level of synthesis, there was not a statistically significant effect of molecule complexity. This implies that molecule complexity may be an important factor in the automated optimization of strains producing those molecules by our pipeline, but it does not significantly impact the low level “proof-of-concept” production of those molecules. The strongest correlation with titer tier was the minimum number of heterologous enzymes required for a molecule’s synthesis. This is not surprising given that for each heterologous enzyme in the pathway introduced into the host organism, there is a probability that either the protein or required reaction chemistry fail to function in the new host, making the probability of successful production drop with each additional heterologous gene required. Despite that, our methods produced multiple hits for pathways containing 8 to 11 heterologous genes.

**Figure 7.**
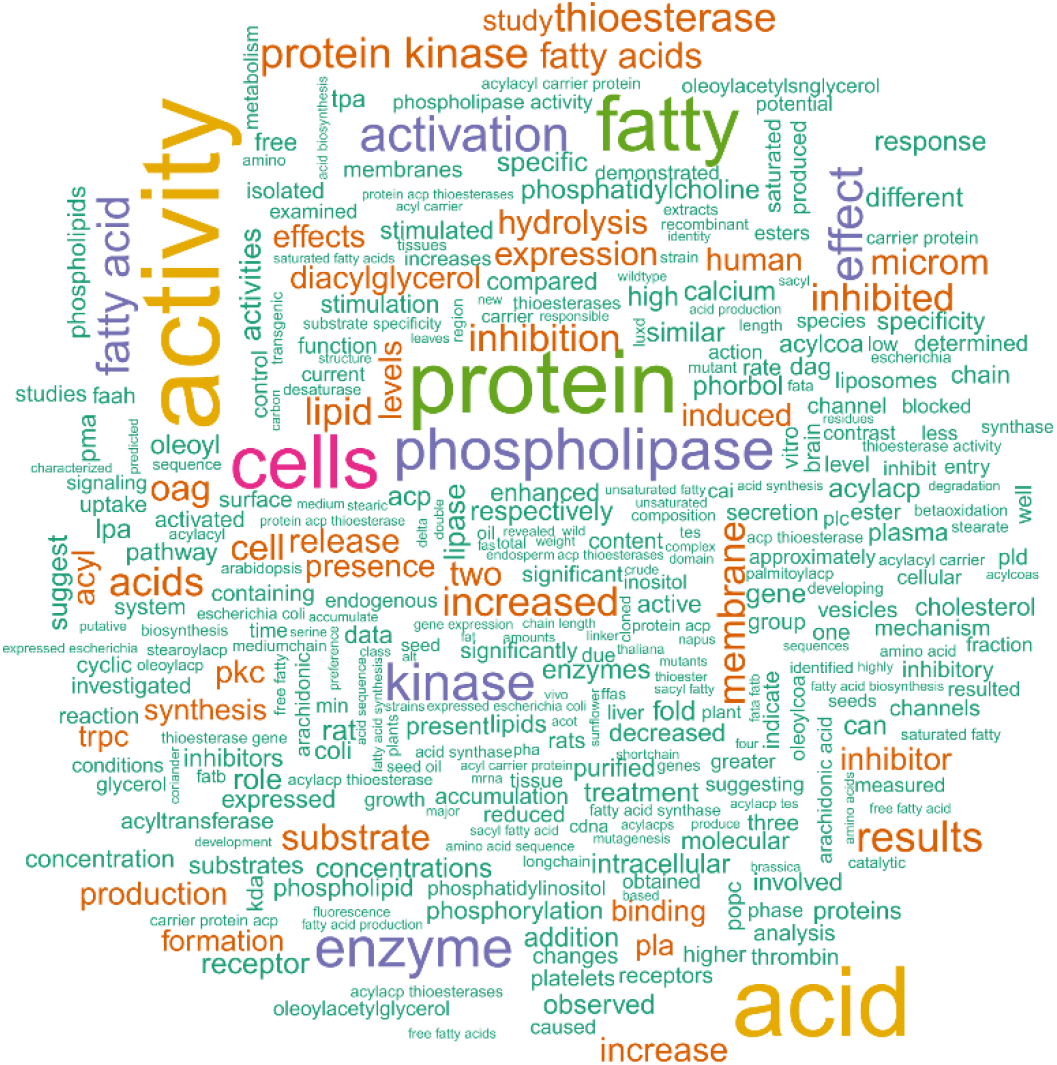
Word-cloud obtained from literature-mining for the enzyme “oleoyl hydrolase”, EC# 3.2.1.14.

**Figure 8.**
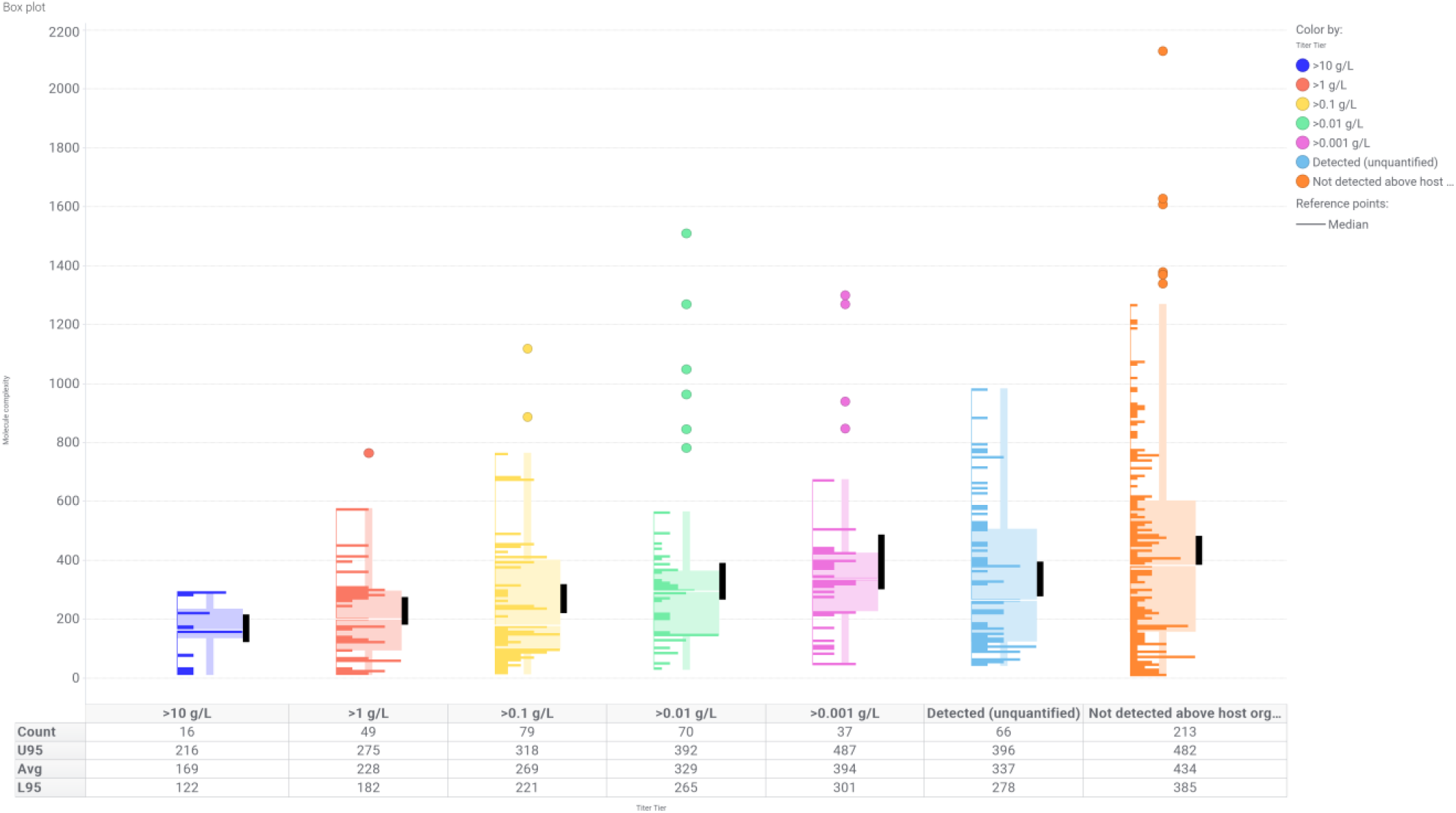
Box-plot of the distribution of molecule complexities at each titer tier. The black bars represent the 95% CI of the mean. The population size (Count), mean (Avg), upper 95% (U95) and lower 95% (L95) confidence interval bounds are listed explicitly in the statistics table below each plot. Data points include both 96-well plate and 2L tank observed titers as separate data points.

**Figure 9.**
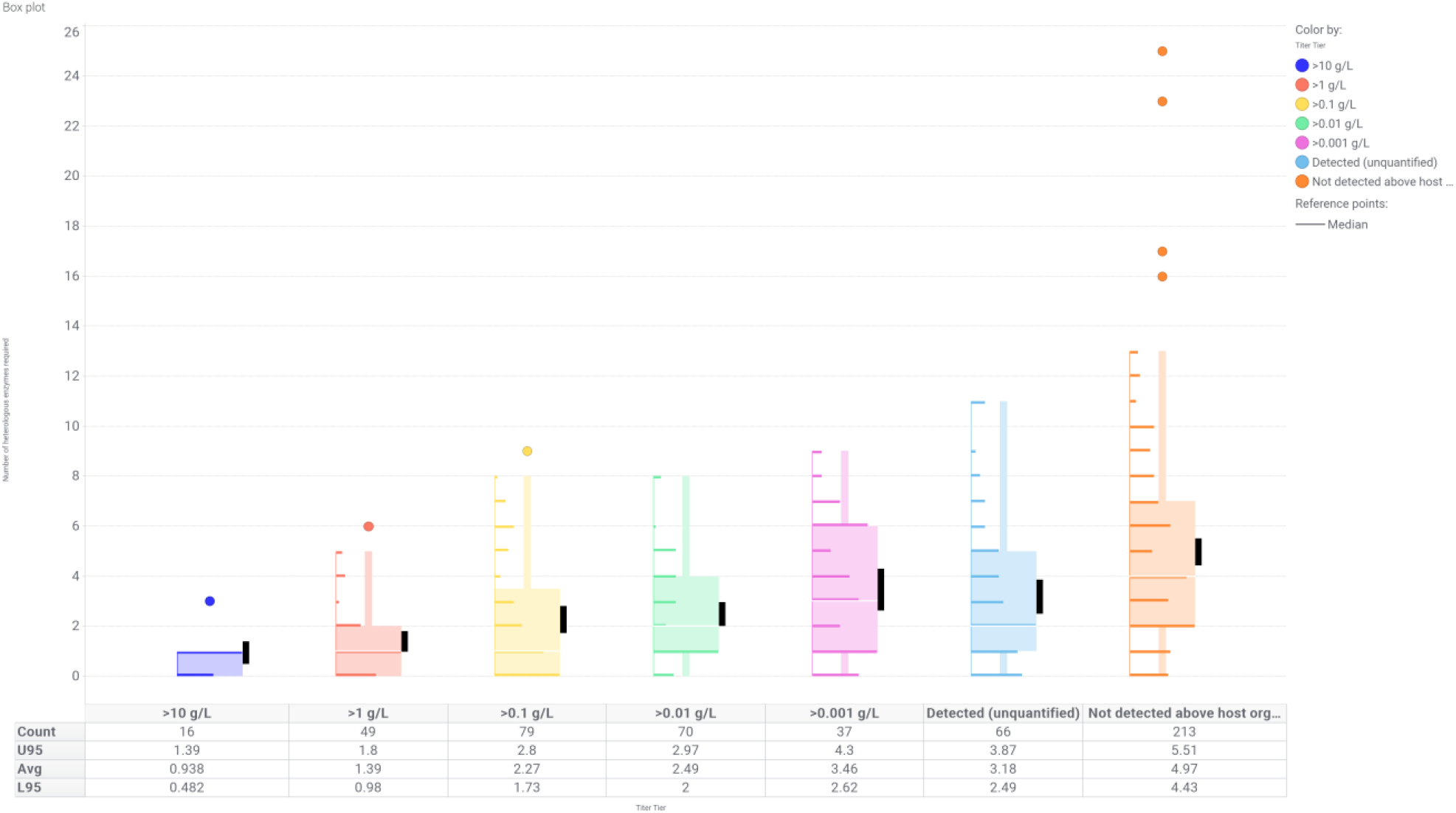
Box-plot of the distributions of the minimum number of heterologous genes required to synthesize the target molecules at each titer tier Note that in some cases, additional heterologous enzymes were added to improve flux above the capabilities of hthe host pathway. The black bars represent the 95% CI of the mean. The population size (Count), mean (Avg), upper 95% (U95) and lower 95% (L95) confidence interval bounds are listed explicitly in the statistics table below each plot. Data points include both 96-well plate and 2L tank observed titers as separate data points.

**Figure 10.**
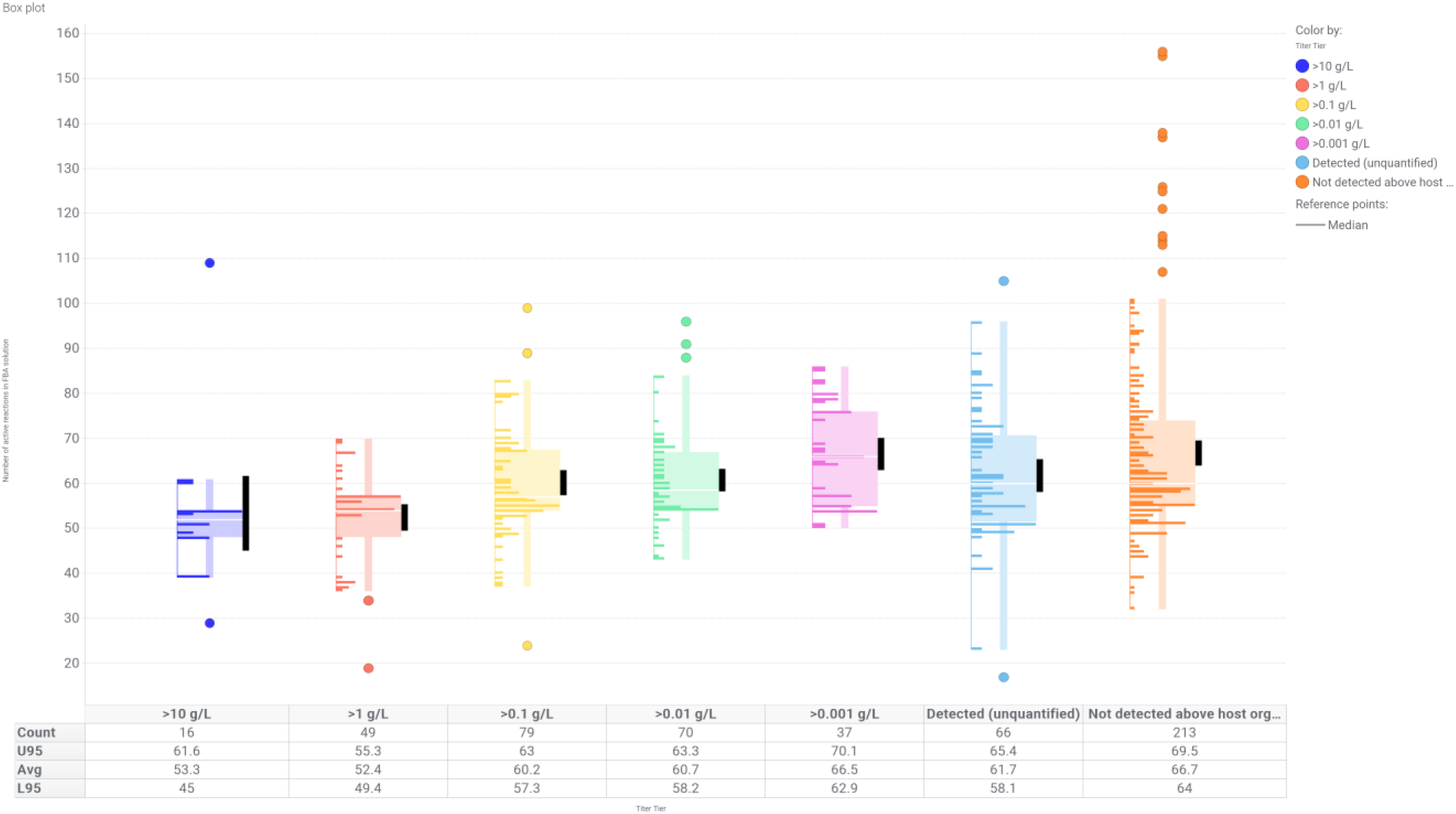
Box-plot of the distributions of the number of active reactions in the maximal yield FBA solution from the Route Finder for the target molecules at each titer tier. The black bars represent the 95% CI of the mean. The population size (Count), mean (Avg), upper 95% (U95) and lower 95% (L95) confidence interval bounds are listed explicitly in the statistics table below each plot. Data points include both 96-well plate and 2L tank observed titers as separate data points.

## Methods

### Route Finder Algorithm

The Route Finder is based on flux-balance analysis (FBA)^56^. Starting with published genome-scale metabolic models for *S. cerevisiae*^43^ and *E. coli*^44^, we extended these models to include all known heterologous reactions using our Universal Set of Reactions. Furthermore, all reversible reaction stoichiometries were split into two unidirectional reactions such that all reaction fluxes **v** were strictly non-negative. We then ran FBA on this extended model **S** with an objective function weight vector **w. w** was defined with negative weights for all reactions. Reactions that were native to the model organism received a small negative weight and heterologous reactions were assigned a ∼100x larger negative weight. This effectively penalized the use of any heterologous reactions unless required to produce the target metabolite. The Route Finder performed FBA on these extended models with COBRApy v0.3.2 to find the minimal set of heterologous reactions that could produce an arbitrarily small demand flux *c* on the target metabolite for overproduction:

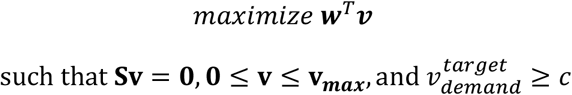

The resulting flux distributions from these models were further analyzed for thermodynamic feasibility, overall stoichiometry, and maximum theoretical yield. Roughly a quarter of the routes were checked manually by Amyris strain engineers, which fed back into successive refinements of Lila’s Route Finder algorithm by altering elements of **w** or **v**_***max***_.

### Enzyme Selection Algorithm

The enzyme selection algorithm filters amino acid sequences using scientific criteria prioritized in the following order: *(i)* the fraction of PubMed abstracts mentioning an enzyme of interest and all its aliases, that also mention the target molecule of interest or a specific host organism, *(ii)* existence of k_m_ values in BRENDA, *(iii)* experimental evidence for the existence of the genetic sequence at the protein or mRNA level from the Uniprot database, *(iv)* phylogenetic tree topology, and *(v)* sequence diversity from a multiple sequence alignment.

If the first pass of the algorithm chooses more variants than can be reasonably synthesized, it attempts to reduce the candidates by making the selection criteria more stringent. If there are too few variants, it attempts to add back candidates by relaxing the selection criteria. It does this iteratively for each enzyme until convergence. In many cases the algorithm starts with over 100 candidate enzymes for a given EC#, and can narrow it down to four candidates within ten iterations (<5s runtime).

The amino acid sequences selected for each EC# are then codon-optimized for expression in the desired host, most frequently at Amyris the *S. cerevisiae* strain CEN.PK2^57^ using Genotype Specification Language^41^ (GSL) compiler with proprietary codon optimization extensions, and stored in the Knowledge Store. To track each gene and anonymize the submission to the gene synthesis vendor, we devised a random/unique hash key for each DNA sequence that tracks the molecule ID, Uniprot ID, EC#, source organism, and versions/parameters of algorithms that were used to generate the DNA sequences.

To decide on these criteria, we conducted roughly a dozen hours of interviews with Amyris strain engineers and binned their responses into a ranked-choice scheme summarized in SI Table 2.

### Towards a Curated Knowledge Store of Molecules, Genes, and Proteins

The goal of building the Knowledge Store was to be able to store various public and in-house facts pertaining to the DBTL cycle, in order to drive automated strain design and re-design. At the outset of this effort, we conducted a gap analysis to assess what data were already routinely stored at Amyris and what additional data were needed to support Lila. Briefly, the following systems were already in place:

- **Strain** database, storing strain genotypes, a record of integrations/deletions/modifications, and all strain lineages in the history of Amyris, together with QC data
- **Part** database, storing the inventory of all DNA parts with their sequence information, and QC data provided by next-generation sequencing
- **High-throughput Screening** database, storing results of dozens of distinct 96-well plate assays in high- and medium-throughput format, together with experimental conditions
- **Fermentation & Analytical Chemistry** databases, storing results of all benchtop & full-scale fermentations, including time series traces of off-gas measurements, analytical chemistry measurements from GC/LC, mass balance calculations, and experimental conditions
- **Omics** databases, storing results of all transcriptomics, metabolomics, and proteomics experiments, along with experimental conditions
- **Sequencing** database, storing results of all whole-genome sequences and analysis therein, such as identification of SNPs, indels, transpositions, and copy number changes

As this was already quite a substantial list of databases covering Build & Test and we were keen not to reinvent the wheel, we focused on building out specific storage to support Design & Analyze (re-design). These include:

- **Molecule** database, recording public & private identifiers for every molecule & metabolite that may ever appear in a biochemical pathway or a chemical synthesis; together with information on molecule name aliases, safety, hazard/precaution, toxicity to human and microbe, and measured & predicted physico-chemical attributes such as boiling point, Kovats retention index, viscosity, partition coefficient
- **Metabolic model** database, storing all publicly known and privately curated biochemical reactions, stoichiometries of participating metabolites, subcellular localization of each metabolite, and mappings between reactions and enzymes
- **Route** database, storing lists of reactions comprising a biochemical pathway to a target molecule, allowing the storage of routes generated by different computational algorithms (flux-balance analysis, shortest-path, manually curated), and affiliated data such as maximum theoretical yield, yield-productivity cost curves, and Gibbs’ free energies of reactions
- **Protein/enzyme/gene/organism** (central dogma) database, storing public identifiers for proteins, enzymes (EC#s), genes, and organisms, starting with the Uniprot, Brenda, Metacyc, and NCBI databases and extending these to include literature-mined and privately curated entities; literature information about enzyme activity & catalytic rates, protein multimers forming an enzyme, taxonomic lineages, mappings between enzymes and proteins, and name aliases for genes, enzymes, and proteins
- **Business-specific** databases, storing in-house information on running the DBTL pipeline such as updated molecule status, contents of a genetic design, mapping between design and DNA parts, details of gene synthesis orders and codon optimization parameters, and details of machine learning algorithms

The stores above are implemented as a Postgres relational database on a dedicated load-balanced set of servers. For data that are retrieved from public sources, we have also written automated daily, weekly, or monthly scripts that synchronize our databases with newly-available public information. The frequency of these updates depends on the frequency with which the public database is updated and the volume of new data to be imported. Every row of every database table is timestamped with create & modify times, allowing us to rapidly re-generate a copy of the database from any arbitrary point in time and recreate the dataset that may have fed a particular algorithm.

There were several technical challenges in creating, populating, and cross-indexing tables in the knowledge store. First, the naming schemes for different molecules, metabolites, and reactions had to be unified. For example, “ethanol” has 4,292 different names in different databases that all indicate C_2_H_5_OH; “muconic acid” has three different isomers and two charge states that needed to be reconciled, and only a specific subset of these isomers/charge states actually participate in biochemical reactions. We solved this problem by assigning each molecule/metabolite a unique numeric ID, and matching this numeric ID to all possible aliases from different databases. To reconcile charge and isomeric states, we used IUPAC International Chemical Identifiers (InChI), whose three-part format allowed us to distinguish between the bond matrix, the isomer, and the charge. Second, the database schemas had to be merged between the knowledge store and Amyris’ existing database of enzymes and metabolic models. Merging the two database schemas involved merging & cross-indexing metabolites with molecules, merging enzymes, and merging reactions. Finally, we also cross-indexed Amyris’s repository of hundreds of thousands of DNA parts with the Knowledge Store’s schema for genes and enzymes. This was a complex effort involving lookups between Amyris’ internal store for gene names, public curated databases for host organisms such as Saccharomyces Genome Database and EcoCyc, and public sequence databases such as Uniref and NCBI RefSeq. Completing this cross-indexing has enabled fast in-house searches of DNA for heterologous enzymes, as well as the ability to rapidly map novel designs back to existing Amyris strains.

### Determining Factor Levels in Combinatorial Designs

We first finalized the list of factors that would be part of the combinatorial design (Table 1). We then worked against the allocated pipeline capacity as well as the base strain design to dial down the number of factors and levels so as to obtain roughly four strains/variants per molecule and to maximize re-use of DNA parts in subsequent cycles. Many factors that we hoped would float freely, such as terminator, were eventually bound by the restrictions of DNA assembly or re-use. Nevertheless, our designs cross promoter, terminator, and gene with locus in several flavors of designed experiments to aid future mapping between genotype and phenotype.

### Strain optimization algorithms using metabolite data

#### Remove all reactions using or producing off-pathway metabolite

In this algorithm we leverage a curated table of reactions to identify all reactions in which the identified off-pathway metabolite is either a substrate or product. These reactions are next mapped to native proteins using our metabolic model. Finally, we created updated strain designs that replace the promoter of these native genes with one or more weak promoters.

#### Remove all single reactions that KO ability to produce off-pathway metabolite

In this algorithm each off-pathway metabolite is first used to create a metabolic flux model capable of generating, *in silico*, that metabolite. Then, each reaction in the original route is removed, and the metabolic flux model is re-run. If removal of a reaction causes the model to be unable to generate the off-pathway metabolite (i.e., the FBA solution fails to converge on a solution), then that reaction is deemed essential for producing the off-pathway metabolite. To eliminate the off-pathway metabolite, we then generated strain designs that knockout (or titrate the promoter to lower expression) these off-pathway genes.

#### Remove all pairs of reactions that KO ability to produce off-pathway metabolite

In some cases, the metabolic model is able to find more than one route to a given off-pathway metabolite, and in these cases no single knockout is sufficient to eliminate accumulation of that unwanted molecule. In these cases, the re-design algorithm identifies pairs of proteins/reactions that when removed together should eliminate production of the unwanted metabolite.

#### Knock out all reactions that parsimoniously link route to off-pathway metabolite

Finally, we recognized that many off-pathway metabolites would be generated by carbon leaking one or two reaction steps from the core pathway, for example when a native enzyme effectively diverts flux away from a step in the engineered pathway. In these cases, we ran our metabolic model to make a route to our off-pathway metabolite using a variant of Flux Balance Analysis in which all reactions in the original design were given at no cost to the objective function. This approach biased flux towards the original pathway and therefore identified possible carbon drains that branched from the original solution. These branches were then selectively downregulated.

Strain designs implementing these algorithms were generated, had DNA parts assembled, passed through a transformation pipeline, and phenotyped. As in the case with proteomics, we implemented a diverse panel of re-design algorithms. We expect the success of each algorithm to be context-dependent, for example varying depending on the off-pathway pool size, the subsystem of metabolism being engineered, or the percentage of theoretical yield of the parent strain, and future machine learning efforts can be focused on classifying when each algorithm is most likely to succeed.

## Discussion

In this paper, we describe the design an implementation of a computational system to automate the entire process of microbial strain engineering and to rapidly scale the biosynthesis of molecules from milligram to kilogram titers with minimal human input. This Automated Scientist we call Lila is implemented as software modules whose functions mimic the cognitive functions of a human scientist: memory, perception, learning, and decision-making. By evolving the Automated Scientist from an expert system^58^ to an empirical learning system, we were able to bootstrap its initial runs without much initial data, and use an iterative software development cycle to add features and refine models and decision-making frameworks as data were gathered.

In its maiden run as an expert system, Lila distilled its design principles from domain experts (i.e., biologists) and articulated them as computational rules. Certain principles were hard coded. For example, we taught Lila that all P450s must be paired with an appropriate cytochrome P450 reductase, and that all non-native metabolic reactions must be catalyzed by a heterologous protein. These rules were generated deductively from an underlying theory of how cellular metabolism is believed to work, and as a result did not depend in any explicit fashion on in-house data such as metabolomics, proteomics, or titer. Although a rules-based approach was exceptionally fruitful at the initial stages, leading to the creation of strains capable of making several hundred different target molecules, it is largely unable to self-correct or learn over time. As a result, the rate of strain improvement would slow and eventually stop unless the Automated Scientist can observe how strain designs fail and automatically take corrective action.

As we collected more data, the Lila software’s ability to diagnose design sub-optimalities and incorporate feedback from domain experts steadily improved. This improvement was evidenced by the 2x, 4x, or 8x improvements in titer over a matter of months after Lila’s maiden run, and the ability of the Designer module to rapidly learn bottlenecks and new design rules based on the decoration of its metabolic models with new mass spectrometry data.

Synthetic biology has established itself as an emerging paradigm for how drug discovery, chemical manufacturing, and elucidation of biological mechanisms will be conducted in the future. To this end, we present a framework of how computational design, machine learning, and human intelligence can be integrated to accelerate strain design and optimization to produce a variety of chemical targets via fermentation. Traditional pathway design and optimization require inputs from multiple sources, and manual intervention at multiple points in the process, thereby increasing the timeline and cost of the entire endeavor. The goal of the Lila software is to accelerate this process and provide a computational framework to remove the strain optimization bottlenecks that scientists encounter in this field. The next step towards achieving this objective is to routinely apply machine learning algorithms to make ever more sophisticated predictions on how genetic changes impact cellular physiology using data generated from all previous experiments.

## Acknowledgements

We thank all Amyris R&D teams that directly supported this work, including Automated Strain Engineering (M. Christie, K. George, D. Hollis, C. Panackia, N. Patel, K. Chahat, T. Perdue, G. Sagala, P. Yeh), High Throughput Screening and Automation (J. Cragg, H. DePaul, C. Elliot, M. Gustincic, G. Hailu, R. Lao, D. Misumi, D. Nadler, A. Navidi, Y. Park), Analytical Chemistry and Operations (N. Agbonkonkon, A. Chassy, K. Clarke, M. Leavell, S. Mainberger, S. Gaucher, A. Thompson, B. Van Deren, G. Wojciechowski), Biology (K. Benjamin, I. Bogorad, S. Borisova, V. Hsiao, A. McGill, J. Walter), Bioinformatics and Software Engineering (C. Dolan, I. Gilmore, B. Hawthorne, J. Ma, S. Zhang), Fermentation Process Development and Operations (L. Chao, S. Do, T. Leaf, J. Leng, D. McAdam, S. Patel, D. Yim), Bioanalytics (P. Jackson, I. Ribeiro, C. Sandoval, T. Scherbart, A. Zawadzka) and Lab Services (N. Fernandez, J. Morata, B. Tanjoco). We also would like to acknowledge J. Ubersax, D. Abbott, M. Leavell for ideation and co-writing the grant that funded this work, and K. Benjamin and M. Leavell for manuscript feedback.

## Supplementary Information

